# 4D force patterning enables spatial control of angiogenesis

**DOI:** 10.1101/2025.11.11.687921

**Authors:** Sina Kheiri, Jessica Shah, Peiyuan Chai, Shashaank A. Venkatesh, Ryan A. Flynn, Roger D. Kamm, Ritu Raman

## Abstract

Engineering organized microvascular networks remains a critical challenge in tissue engineering and regenerative medicine. While biochemical approaches for patterning angiogenesis via growth factor delivery have shown promise, their inability to pattern sustained growth factors with spatiotemporal control limits effectiveness. Here, we demonstrate that dynamically patterned mechanical forces enable precise spatiotemporal control over angiogenic sprouting. We developed a magnetically actuated human vessel-on-a-chip platform that integrates a perfusable endothelialized microchannel within a collagen matrix and allows non-invasive and tunable mechanical stimulation across three spatial dimensions and time (4D). Using an automated 3-axis actuator, we systematically investigated how strain magnitude, frequency, and direction modulate endothelial cell behavior and vessel morphogenesis. Dynamic mechanical stimulation at physiological strain magnitudes (5–15%) enhanced endothelial alignment and barrier function while promoting angiogenesis in a strain-magnitude–dependent manner: lower dynamic strain (5%) maximized sprout initiation, whereas higher dynamic strain (15%) promoted elongation of sprouts. Sequential reorientation of strain direction reprogrammed sprouting trajectories along X, Y, and Z directions, generating complex sprout geometries such as L-shaped branches. RNA sequencing revealed mechanically induced transcriptional profiles distinct from unstimulated controls, characterized by upregulation of genes associated with angiogenesis, mechanotransduction, and extracellular matrix remodeling. Functional perturbation of Piezo1 reduced strain-induced sprouting without altering barrier stabilization, indicating that dynamic mechanical stimulation engages multiple mechanotransduction pathways to regulate angiogenesis. Collectively, these findings establish a strategy for spatiotemporally controlled angiogenesis through 4D force patterning to program vascular morphogenesis while preserving function. This approach provides a foundation for engineering hierarchically organized vascular networks for tissue regeneration.

**Significance:** Generation of spatially organized, perfusable microvascular networks is essential for building functional human tissues. Biochemical approaches to pattern angiogenesis rely on diffusive growth factors, which limit control over spatiotemporal sprouting dynamics. Here, we demonstrate that dynamically patterned mechanical forces direct vascular morphogenesis across three spatial dimensions and time (4D). Using a magnetically actuated human vessel-on-a-chip, we show how strain magnitude and orientation govern angiogenic sprouting and reveal transcriptional programs linking mechanical cues to observed functional changes. For the first time, we show that dynamic reorientation of imposed forces can reprogram angiogenic trajectories in real-time. This platform enables programmable mechanical control of angiogenesis and systematic dissection of mechanotransduction pathways, advancing strategies for tissue vascularization, and modeling mechanically regulated vascular diseases.

## Introduction

The vascular network spans a wide range of length scales, from millimeter- to centimeter-scale arteries and veins to microscale capillaries about 5–10 µm in diameter, enabling the transport of oxygen and nutrients while removing metabolic waste.(1) Perfusable vasculature is thus essential for the integration of engineered tissues, as diffusion limits cell viability to only a few hundred micrometers, leading to hypoxia, necrosis, and functional loss.(2, 3) Yet, replicating the intricate structure and function of native microvasculature remains a significant challenges in tissue engineering.(1) Although significant progress has been made in engineering microvascular networks within tissues, current approaches generally rely on two strategies: self-organization or pre-patterning.(4, 5) Self-organization of endothelial cells embedded within hydrogels produces microscale networks with natural topology, but suffers from random connectivity and limited control over network geometry. In contrast, pre-patterning of vascular templates via methods like 3D bioprinting provides precise spatial control but is constrained by resolution limitations (typically 100s of µm), preventing the recapitulation of fine-scale vascular hierarchies and spontaneous remodeling.(6, 7) Consequently, fabrication methods that enable hierarchically organized, precisely patterned, and perfusable microvasculature hold great promise for advancing engineered tissue models for *in vitro* disease modeling and *in vivo* regenerative medicine applications.

Biochemical patterning strategies have been extensively employed to induce vascular formation by stimulating angiogenesis through pro-angiogenic cues, primarily vascular endothelial growth factor (VEGF).(8–10) Approaches such as functionalized bioinks for controlled VEGF release and aptamer-tethered VEGF photopatterning have enabled partial spatial regulation of sprouting and network formation.(11, 12) Despite these advances, biochemical methods remain limited by the rapid diffusion and short half-life of VEGF, which hinder stable, localized gradients in three-dimensional (3D) environments. Furthermore, because angiogenesis is an inherently spatiotemporal process involving phases of growth, regression, and remodeling, static or one-time VEGF delivery often results in irregular and disorganized vascular networks that deviate from native microvascular architecture.(13)

In native tissues, angiogenesis is governed not only by biochemical cues but also by biomechanical signals (14). Endothelial cells are highly mechanosensitive and continuously exposed to both fluid flow-induced mechanical stimuli (shear stress and cyclic pressure generated by blood flow) and extracellular matrix (ECM) deformation-induced mechanical stimuli (tension and compression generated by tissue movement surrounding vessels) which collectively influence sprouting, alignment, and barrier function.(15) *In vivo,* blood vessels undergo dynamic mechanical stretching from tissue growth and movement, which influence the spatial patterns and orientation of vascular networks, yet these forces remain largely understudied compared to hemodynamic forces.(16) Early work demonstrated that endothelial spheroids cultured on statically stretched collagen gels exhibited directional outgrowth along the axis of strain (17). While this showed that strain can bias sprouting direction, placing spheroids on top of stretched gel limited direct force transmission to the cells and lacked physiological cell-matrix embedding. To directly apply mechanical forces in the same plane as the cells, later systems delivered strain to 2D cell monolayers via deformable 2D substrates, revealing that mechanical stretch influences network elongation (18), cell and cytoskeletal alignment (19), and reorientation in response to changing strain direction.(20) While these studies provided key insights into how 2D endothelial monolayers respond to mechanical cues, they did not fully capture the complex spatial architecture, cell-matrix interactions, and mechanical gradients that influence endothelial behavior in native 3D environments.

Recent efforts have begun to address this limitation by incorporating mechanical stimulation into 3D vessel-on-a-chip platforms. In one approach, 3D endothelialized lumens embedded in fibrin hydrogels were cyclically stretched by deflecting a thin PDMS membrane using negative pressure (21). While this method enabled mechanical stimulation of 3D tissue, it lacked precise spatial control of force patterning. Another approach applied strain to endothelial cell-lined microchannels by dynamically stretching a gelatin-based hydrogel perpendicular to the channel axis (22). Although this platform allows for more controlled directional deformation, the hydrogel formulation was optimized for mechanical extensibility rather than biological remodeling, limiting its ability to support angiogenic sprouting. More recently, a PDMS-based stretchable device applied static mechanical stretching to an engineered vessel, demonstrating that strain direction significantly influences sprouting behavior (23). Strain perpendicular to the vessel promoted angiogenic outgrowth and alignment, while strain parallel to the vessel inhibited sprouting, highlighting the importance of directional mechanical cues in angiogenesis. While these platforms expanded the ability to study mechanical stimulation in 3D vascular tissues, the applied forces remain global and are typically constrained to one or two axes, limiting the ability to deliver spatially patterned, multi-axial strain with high spatiotemporal precision. As such, key questions remain about how multi-axial and dynamically varying mechanical cues applied along the x, y, and z axes regulate angiogenic sprouting and barrier function in 3D tissues. A central challenge in understanding and directing vascular patterning is thus the difficulty of delivering tunable, non-invasive mechanical stimulation in both 3D space and time (4D) to mimic evolving *in vivo* mechanical environments.

To address limitations of conventional mechanical stimulation methods, magnetic stimulation offers a non-invasive means to apply controlled mechanical forces in engineered tissues.(24, 25) We have previously established a magnetic matrix actuation (MagMA) platform in which magnetic actuators embedded in an ECM can be non-invasively actuated by an external magnet, enabling precise spatial patterning of forces within an engineered tissue (26, 27). Here, we adapted MagMA to enable 3D actuation of a collagen hydrogel containing a 3D perfusable vascular channel lined with human endothelial cells. To deliver dynamic, multi-axial strain in 4D, we used a motorized 3-axis actuator to control the position, height, and speed of external magnets above the device, enabling spatial and temporal control over strain magnitude, frequency, and direction within the tissue. By tuning strain magnitude, we observed distinct effects on vascular morphology: lower strain increased sprout number, while higher strain promoted elongation of fewer sprouts. Beyond enhancing sprout initiation and elongation along a single axis, our platform enabled sequential multi-directional mechanical stimulation to dynamically reorient 3D sprouting trajectories along X, Y, and Z dimensions over time. This approach revealed that endothelial sprouts could adapt their growth trajectories in response to changing mechanical inputs, generating complex geometries such as L-shaped sprouts. Transcriptomic profiling further demonstrated that dynamic strain activated angiogenic and mechanosensitive gene programs while preserving barrier function. Genetic perturbation of Piezo1 attenuated strain-induced sprouting while preserving barrier stabilization, suggesting a role for specific force-sensing pathways in mechanically regulated angiogenesis. Together, these findings establish a platform for mechanical stimulation of a human vessel-on-a-chip, highlighting the potential for spatiotemporally patterned forces to program 4D morphogenesis of microvasculature in engineered tissues.

## Results

### Design and Fabrication of Magnetically Actuated Human Vessel-on-a-Chip Platform

A human vessel-on-a-chip system was engineered to impose controlled, dynamic mechanical strain on an endothelialized microchannel through an integrated magnetic actuation mechanism (Fig. 1A; Fig. S1A). The device was fabricated using a multi-step molding process (***see Materials and Methods***; Fig. S1B; Movie S1). Briefly, a glass rod was inserted into the device mold to define a central lumen within a type I collagen matrix (Fig. 1A, steps 1–2). To ensure reproducible positioning of the magnetic actuator relative to the engineered vessel, a custom 3D-printed stamp containing a pin that interfaces with a corresponding slot in the acrylic device was used to pattern a dedicated actuator cavity in the gel (Fig. 1A, steps 3-4; Fig. S1C). The actuator was embedded within this predefined cavity at an edge-to-edge distance of 535 ± 20 µm prior to encapsulation with an additional collagen layer (Fig. 1A, steps 5-6; Fig. S1D). Following device assembly, human umbilical vein endothelial cells (HUVECs) were seeded into the channel (Fig. 1A, step 7), forming a confluent endothelium within 48 h. The resulting endothelialized microchannel formed a lumenized vessel, providing a reproducible platform to investigate the effects of controlled mechanical stimulation on angiogenesis. (Fig. 1A, step 8).

**Figure 1.**
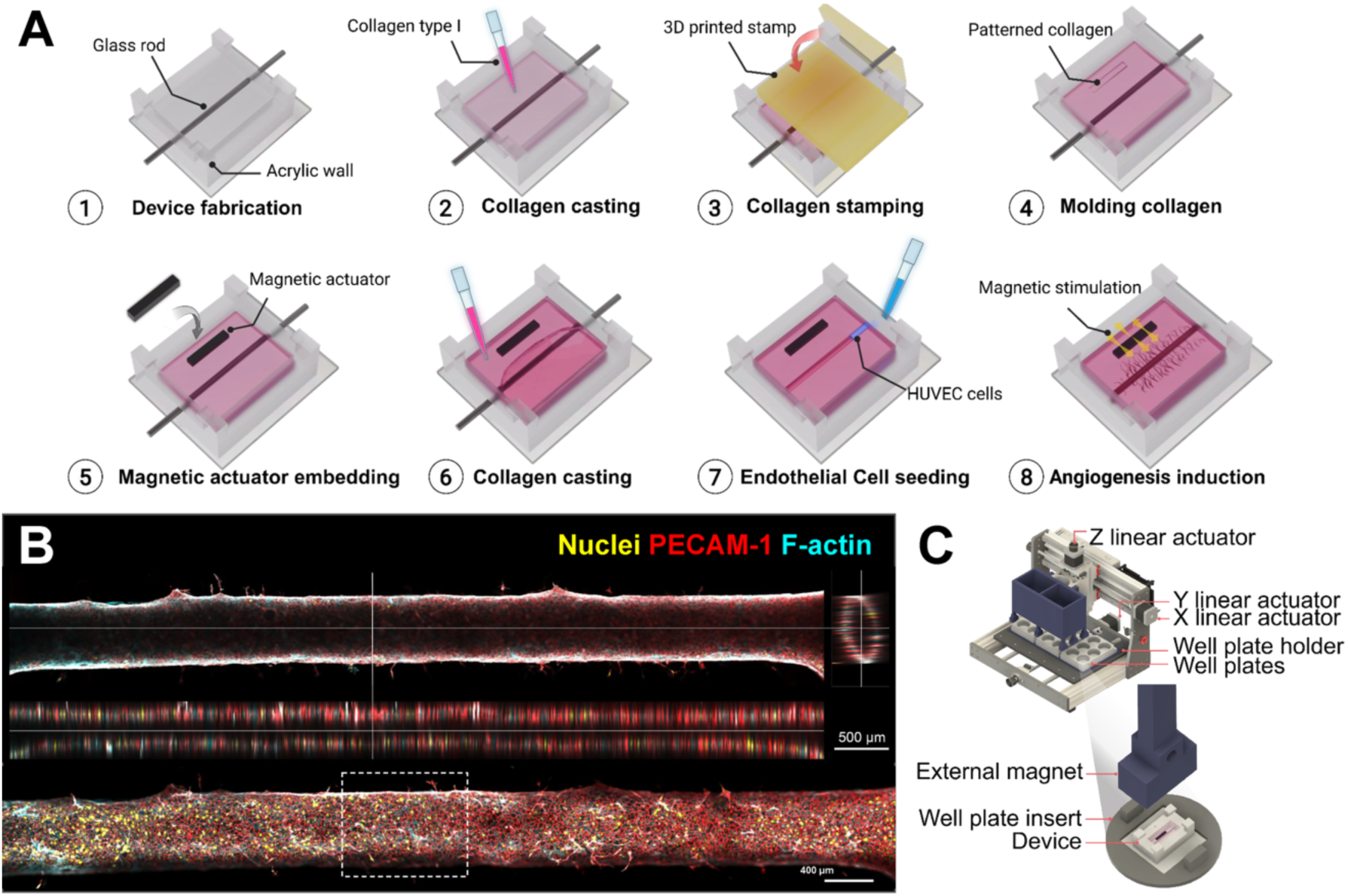
Development of a magnetically actuated vessel-on-chip for mechanical modulation of angiogenesis. (*A*) Schematic workflow of magnetic actuator integration into the vessel-on-a-chip. A glass rod was inserted to define a hollow channel within the collagen matrix (steps 1–2). A 3D-printed stamp patterned a slot for reproducible positioning of the magnetic actuator (steps 3–4). The magnetic actuator was embedded adjacent to the lumen and encapsulated with an additional collagen layer (steps 5–6). HUVECs were seeded into the channel to form a confluent endothelium within 48 h (step 7). Controlled dynamical stimulation was then applied using an external permanent magnet (step 8). (*B*) Representative confocal microscopy images of the unstimulated engineered vessel. Nuclei (yellow), PECAM-1 (red), and F-actin (cyan). Vertical and horizontal cross-sections demonstrate the hollow lumen structure and complete endothelial lining of the vessel. Scale bar, 500 μm. Stitched z-stack maximum projection shows the full 3D vessel structure. Scale bar, 400 μm. (*C*) Schematic of the automated magnetic actuation platform. The vessel-on-a-chip device is housed within a custom well-plate insert positioned beneath an array of external permanent magnets. Magnet motion is controlled by linear actuators enabling programmable, multidirectional magnetic stimulation across multiple wells for high-throughput mechanical testing.

Confocal fluorescence imaging of PECAM-1 (cell–cell junctions) and F-actin (cytoskeleton) showed that HUVECs formed a circular, perfusable lumen with a diameter of 604 ± 42.2 µm. After 3 days of culture in media, endothelial sprouts extended from the parent vessel into the surrounding collagen matrix, demonstrating that the platform supports spontaneous angiogenic initiation (Fig. 1B). To impose controlled mechanical strain on the formed vessels, a custom magnetic stimulation system was developed to control actuation of the collagen hydrogel within the device. The system was designed for high-throughput stimulation, with vessel-on-a-chip devices mounted in standard 6-well plates using a custom insert that accommodated up to 3 well plates, or 18 chips in total (Fig. 1C). To dynamically control the movement of the composite magnetic matrix, a custom 3-axis actuator was constructed to translate a permanent magnet above the devices with controllable distance, speed, direction, and frequency (***see Materials and Methods***). This movement induced both dynamic and static actuation of the embedded magnetic actuator and surrounding collagen matrix, generating uniform strain along the vessel wall (Movie S2). The strain magnitude was tuned by adjusting the magnetic field strength (Fig. S2A).

Quantitative movement tracking analysis of fluorescent beads embedded in the collagen hydrogel at different locations within the chip (Fig. S2B-C) confirmed that increasing the magnetic field from 250 G to 1,800 G modulated the strain magnitude (Fig. S2A). Strain dynamics were first quantified across four lateral rows to characterize temporal profiles during actuation (Fig. S2B). To more precisely resolve mechanical stimulation at the vessel boundary, strain was quantified across six lateral rows (Fig. S2C). Although a lateral strain gradient was observed within the bulk collagen matrix, strain magnitude at the vessel wall was comparable between sides at both 250 G and 1,800 G. Strain magnitudes of approximately 5% and 15% at the vessel wall were selected as representative physiological conditions (16) to examine the effects of dynamic, spatiotemporally patterned strain on angiogenesis.

To evaluate the impact of mechanical stimulation, constructs were stimulated at 5% and 15% strain magnitude, and the role of dynamic loading was examined by applying either dynamic strain at 1 Hz or static strain (Fig. 2A). Constructs were cultured for 3 days to allow vessel formation prior to stimulation (Fig. S3A). Dynamic mechanical loading was then applied once daily for 1 h for 3 additional days. This stimulation window was selected based on time-course screening across 9 days of stimulation, which identified days 0–3 of stimulation as the phase of most pronounced sprouting response under repeated strain. (Fig. S3B, Movie S3) The 1 h per day mechanical stimulation period was chosen to approximate the intermittent loading experienced during in vivo studies (∼1–2 h) and to avoid the growth suppression reported under prolonged continuous cyclic strain *in vitro*.(14, 28, 29) Mechanical stimulation did not significantly alter the mean sprout diameter (Fig. S4C) but broadened the diameter distribution, increasing the proportion of thicker sprouts (>10 µm) compared to control (Fig. S4D). Unlike many 3D bioprinting approaches that generate vascular channels hundreds of micrometers in diameter, the sprouts formed in our platform remain within a capillary-scale range (30).

**Figure 2.**
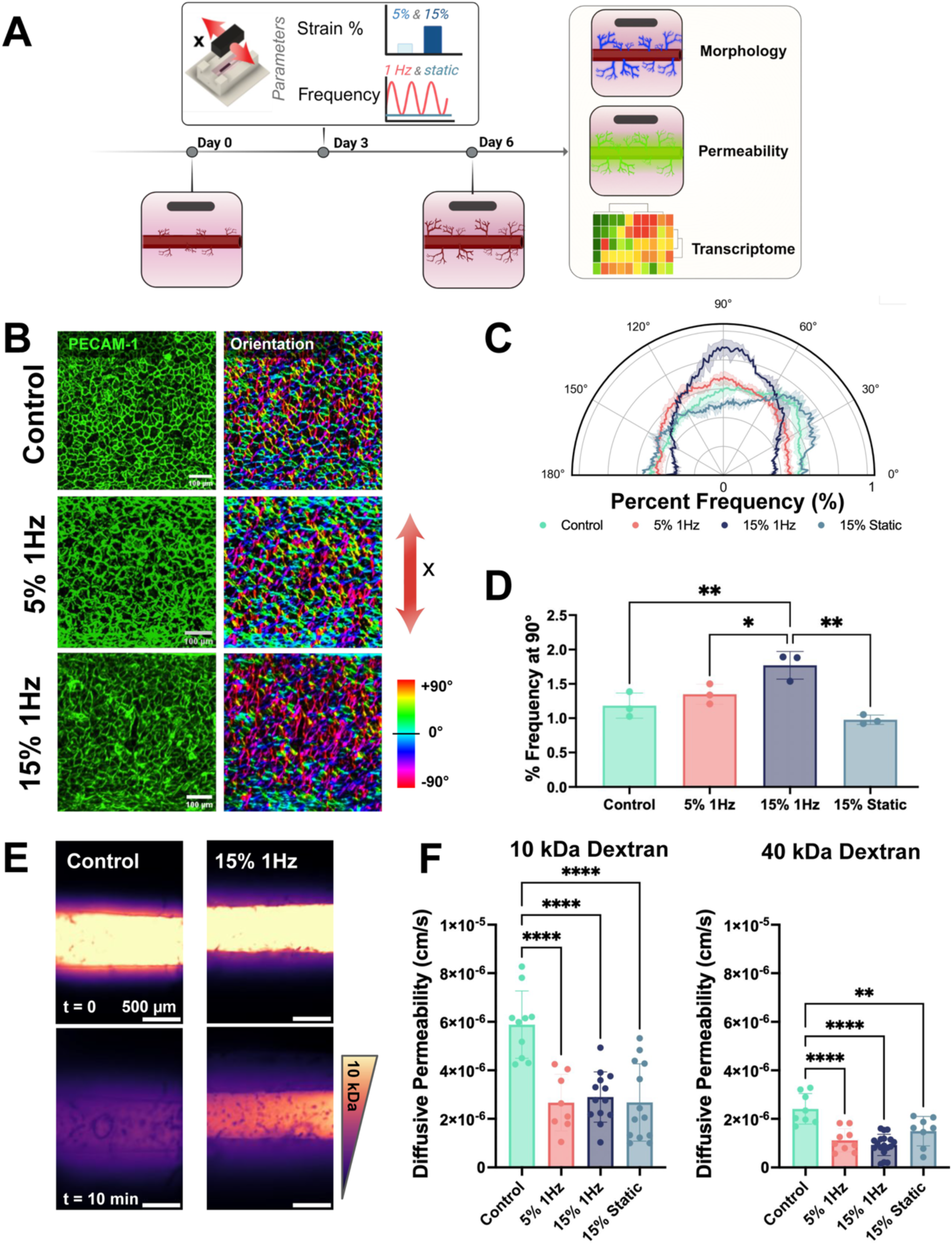
Dynamic mechanical stimulation modulates endothelial cell alignment and barrier properties. (*A*) Experimental timeline showing dynamic or static mechanical stimulation (5% or 15%) of engineered vessels starting on day 3. Vessel morphology, permeability, and transcriptomic profiling were assessed after 3 days of stimulation. (*B*) Representative confocal images showing PECAM-1 (green) and endothelial orientation maps under control, 5% 1Hz, and 15% 1Hz strain conditions in the parent vessel. Orientation analysis was performed using *OrientationJ*, with color indicating alignment relative to the strain axis. Scale bar, 100 μm. (*C*) Polar plots showing the percentage of cells oriented at each angle relative to the strain axis, quantified using OrientationJ. Plots correspond to control, 5% dynamic, 15% dynamic, and 15% static strain conditions. Shaded regions indicate standard deviation across biological replicates (n=3 independent devices, one ROI per device). (*D*) Quantification of cell alignment frequency at 90°, corresponding to the direction of applied strain. (^ns^p > 0.05, *p < 0.05, **p < 0.01, one-way ANOVA, n=3 independent devices, one ROI per device) (*E*) Representative fluorescence images showing diffusion of a fluorescent tracer (10kDa Dextran) across the endothelial barrier at t = 0 (top row) and t = 10 min (bottom row). Scale bar, 500 μm. (*F*) Quantification of vessel permeability for 10 kDa and 40 kDa dextran tracers under unstimiulated (control) and stimulated conditions. (^ns^p > 0.05, *p < 0.05, **p < 0.01, ***p < 0.001, ****p < 0.0001, one-way ANOVA, n≥8 independent devices).

To isolate the specific effects of mechanical stimulation on angiogenesis in our platform, we evaluated two potential confounding stimuli: exogenous VEGF in the culture medium and magnetic field exposure. We first studied the effect of exogenous VEGF by comparing total sprouting length in vessels cultured in VEGF-free endothelial growth medium versus full culture medium supplemented with 5 ng/mL VEGF. This represents a relatively low VEGF concentration compared to prior microvessel sprouting studies, which often employ higher VEGF doses (e.g., 25-50 ng/mL) or combinatorial growth factor cocktails to induce angiogenesis (31–33). Dynamic strain significantly increased total sprout length with no significant difference between the VEGF-free and VEGF-supplemented groups (Fig. S3C), indicating that exogenous VEGF supplementation is not required for strain-induced angiogenesis in our platform. We next assessed the effect of external magnetic fields alone on angiogenesis by comparing sprouting and barrier function in vessels exposed to magnetic stimulation in the presence or absence of the magnetic actuator. The magnetic field alone, which is substantially weaker than field strengths previously reported to modulate angiogenesis (34), did not significantly affect angiogenic sprouting or barrier function in our platform (Fig. S3D–E). Together, these results indicate that the observed changes in sprouting and barrier function are driven primarily by dynamic mechanical stimulation.

After confirming that exogenous VEGF in the culture medium and magnetic field exposure did not significantly affect angiogenic sprouting or barrier function in our platform, we characterized the impact of mechanical stimulation on vascular morphology and function by measuring cell–cell junction orientation, barrier function (permeability), sprout magnitude and direction, and gene expression via RNA sequencing.

### Strain magnitude and frequency regulate endothelial cell-cell junctions, cell orientation, and barrier function

Endothelial cell organization was examined using PECAM-1 (cell-cell junction) immunostaining and assessed with analysis of cell–cell junction alignment within the parent vessel. Application of 15% dynamic strain resulted in the highest degree of endothelial cell alignment along the axis of deformation compared to 15% static strain, 5% dynamic strain, and unstimulated control conditions (Fig. 2B-D). Orientation analysis demonstrated that dynamic strain enhanced endothelial alignment in a magnitude-dependent manner (Fig. 2C). Cell–cell junction orientation showed a narrower angular distribution under 15% 1Hz strain compared to 5% strain and no-stimulation controls, indicating a higher degree of directional organization. The percentage of cells oriented at 90° increased significantly under 15% dynamic strain relative to both 5% strain and static culture (Fig. 2D). Notably, under static 15% strain conditions, no significant change in junction orientation as compared to unstimulated controls was observed, suggesting that alignment is specifically induced by dynamic mechanical loading in our platform. These findings indicate that dynamic strain promotes directional endothelial organization, with strain magnitude serving as a key regulator of network alignment.

Barrier function of the vessels was evaluated by measuring the diffusion of fluorescent 10 kDa and 40 kDa dextran tracers across the vessel wall. In unstimulated controls, timelapse confocal imaging showed rapid diffusion of dextran into the surrounding hydrogel, whereas vessels subjected to dynamic strain retained fluorescence within the lumen, indicating reduced permeability (Fig. 2E). Quantitative analysis revealed a significant decrease in diffusive permeability for both 10 kDa and 40 kDa dextran in mechanically stimulated tissues. For the 10 kDa tracer, permeability decreased by nearly two-fold across all stimulated conditions, while the 40 kDa tracer showed approximately a 1.5-fold reduction across all stimulated conditions (Fig. 2F).

### Strain magnitude controls angiogenic sprouting direction and magnitude

Mechanical stimulation induced distinct changes in sprouting architecture, as indicated by the merged immunofluorescence images of nuclei, PECAM-1, and F-actin (Fig. 3A). In unstimulated controls, endothelial sprouts appeared short and randomly oriented, whereas dynamic strain at 5% and 15% showed more elongated sprouts projecting outward from the parent vessel. To quantify changes in angiogenic sprouting under different mechanical stimulation conditions, we measured total sprouting length, maximum sprout length, and sprout number. Mechanical stimulation, independent of strain magnitude or whether the loading was static or dynamic, resulted in a substantial increase in total sprouting length compared with unstimulated controls (Fig. 3B). Further analysis of individual sprout morphology showed that the longest sprouts formed specifically under 15% dynamic strain, indicating that higher dynamic strain preferentially promotes sprout elongation (Fig. 3C). In contrast, the 5% dynamic condition yielded the highest number and density of sprouts, demonstrating that lower amplitude dynamic strain most effectively drives sprout initiation (Fig. 3D). While all forms of mechanical stimulation enhanced angiogenesis, the strain regimen determined whether vessel formation occurred through increased sprout initiation or elongation of individual sprouts.

**Figure 3.**
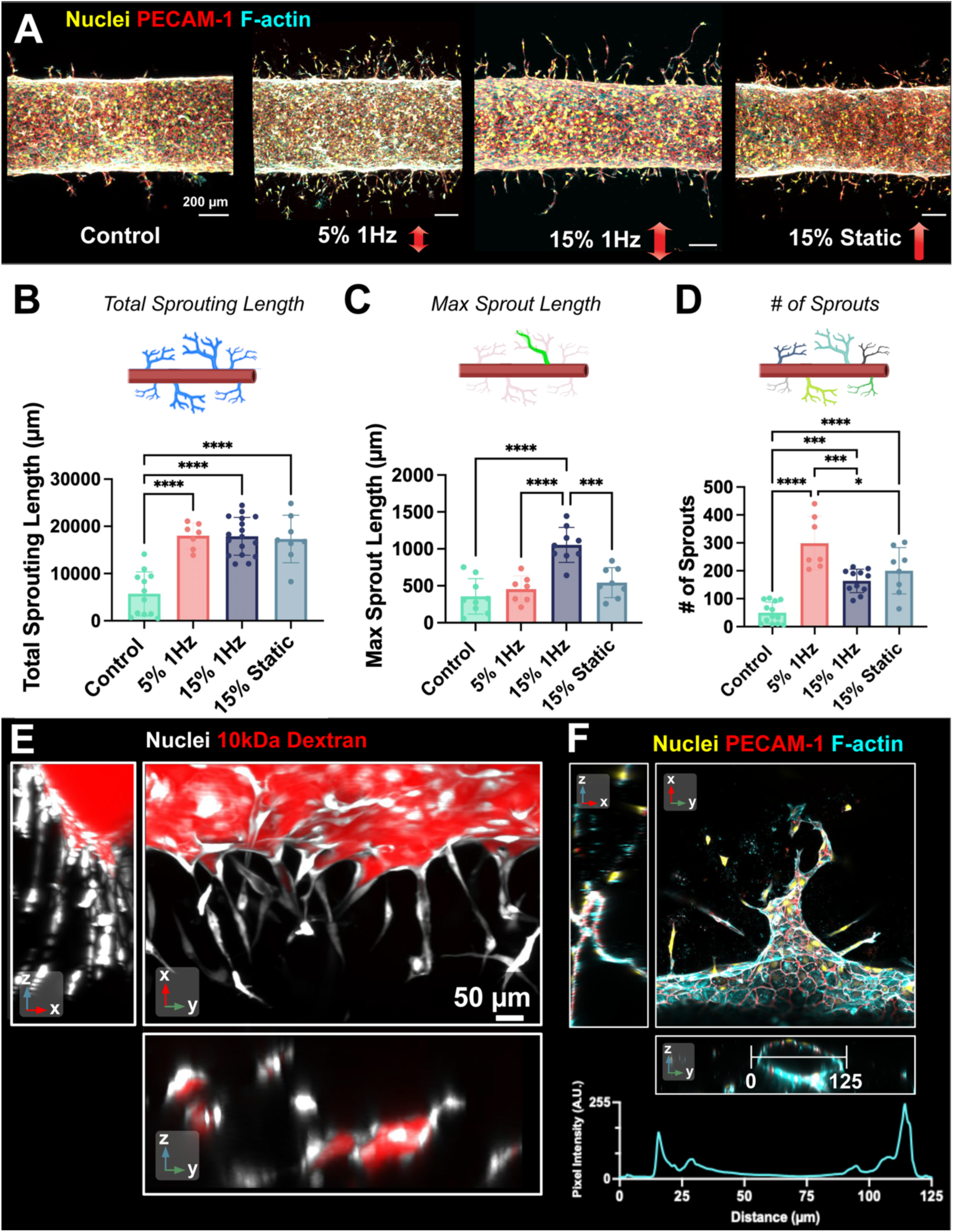
Mechanical strain drives angiogenic sprouting with lumen formation. (*A*) Representative confocal images of engineered vessels stained for nuclei (yellow), PECAM-1 (red), and F-actin (cyan) under control (unstimulated), 5% 1Hz, 15% 1Hz, and 15% static strain conditions. Scale bar, 200 μm. (*B*) Quantification of total sprouting length from the parent vessel under control (unstimulated) and strained conditions. (^ns^p > 0.05, *p < 0.05, **p < 0.01, ***p < 0.001, ****p < 0.0001, one-way ANOVA, n≥7 independent devices, quantified across the entire device) (*C*) Quantification of maximum individual sprout length from the parent vessel under control (unstimulated) and strained conditions. (^ns^p > 0.05, *p < 0.05, **p < 0.01, ***p < 0.001, ****p < 0.0001, one-way ANOVA, n≥7 independent devices, quantified across the entire device) (*D*) Quantification of the number of angiogenic sprouts extending from the parent vessel. (^ns^p > 0.05, *p < 0.05, **p < 0.01, ***p < 0.001, ****p < 0.0001, one-way ANOVA, n≥7 independent devices, quantified across the entire device) (*E*) Confocal images showing diffusion of 10 kDa fluorescent dextran (red) from the parent vessel into the sprouts. Extended orthogonal cross-sections through the full volume confirm dextran localization relative to endothelial nuclei (white). Scale bar, 50 μm.(*F*) Confocal images stained for nuclei (yellow), PECAM-1 (red), and F-actin (cyan) showing lumenized multicellular sprout with continuous endothelial junctions and organized cytoskeletal architecture. Orthogonal projections and line-scan analysis confirm luminal structure. Scale bar, 50 μm

Given that dynamic stimulation produced more pronounced angiogenic responses, we next examined the structural and functional characteristics of the resulting sprouts. We first assessed perfusability by introducing fluorescently tagged 10 kDa dextran into the parent vessel and imaging dye distribution using confocal microscopy (Fig. S4A). The sprouts formed lumenized extensions that were continuous with the parent lumen and capable of supporting dextran transport (Fig. 3E; Movies S4–S5). Consistent with the expected progression of angiogenic sprouting, lumenization was most prominently observed near the base of sprouts adjacent to the parent vessel lumen. More distal regions often remained non-lumenized, reflecting tip-cell–led extension during early stages of sprout outgrowth prior to lumen formation along the sprout stalk. To further characterize lumen formation, we quantified the F-actin intensity profile across individual sprouts (Fig. 3F). This analysis revealed a peripheral actin-rich cortex with a reduced central signal, consistent with a hollow lumen architecture (Movie S6). Together, these findings indicate that dynamic mechanical stimulation promotes the formation of functional, multicellular vascular sprouts rather than partially detached or single-cell protrusions.

To assess whether mechanical strain directs angiogenic growth along its axis, we analyzed spatial differences in sprouting across the left–right (x-axis, along the direction of applied strain) and top–bottom (z-axis, orthogonal to the direction of applied strain) regions of the engineered vessel. Confocal z-stacks were reconstructed in 3D, and a filament tracing algorithm was applied to segment individual sprouts and extract their spatial coordinates (Fig. S5A). Each sprout was assigned to a spatial quadrant (left, right, top, or bottom) by dividing the vessel cross-section into four regions using angular boundaries at 45°, 135°, 225°, and 315°. Sprout coordinates were converted to angular positions to generate polar plots for a representative region of interest (one independent device per condition), where each point represents an individual sprout. Polar plots revealed that control vessels exhibited short, broadly distributed sprouts, whereas 5% strain increased sprouting density along the x-axis, and 15% strain led to longer sprout formation along the x-axis (Fig. 4A). Consistent with these visual trends, quadrant-based analysis showed that unstimulated controls exhibited sprouting across all regions, with a slight baseline enrichment across the x-axis that did not reach statistical significance between left–right and top–bottom regions (p > 0.05) (Fig. 4B). When strain was applied along the x-axis, directional bias along the actuation axis significantly increased, with greater vessel growth observed in the left-right regions (highlighted in blue). At 5% 1Hz strain, sprouting along the strain axis increased 3.0-fold relative to unstimulated controls and 3.1-fold relative to the orthogonal axis (p < 0.0001). At 15% 1Hz strain, sprouting increased 2.4-fold relative to controls and 3.9-fold relative to the orthogonal axis (p < 0.0001). These quadrant-based differences reflect how strain magnitude influences the density of sprout initiation along the applied axis, demonstrating that strain magnitude and direction modulate both the spatial patterning and length of angiogenic sprouts.

**Figure 4.**
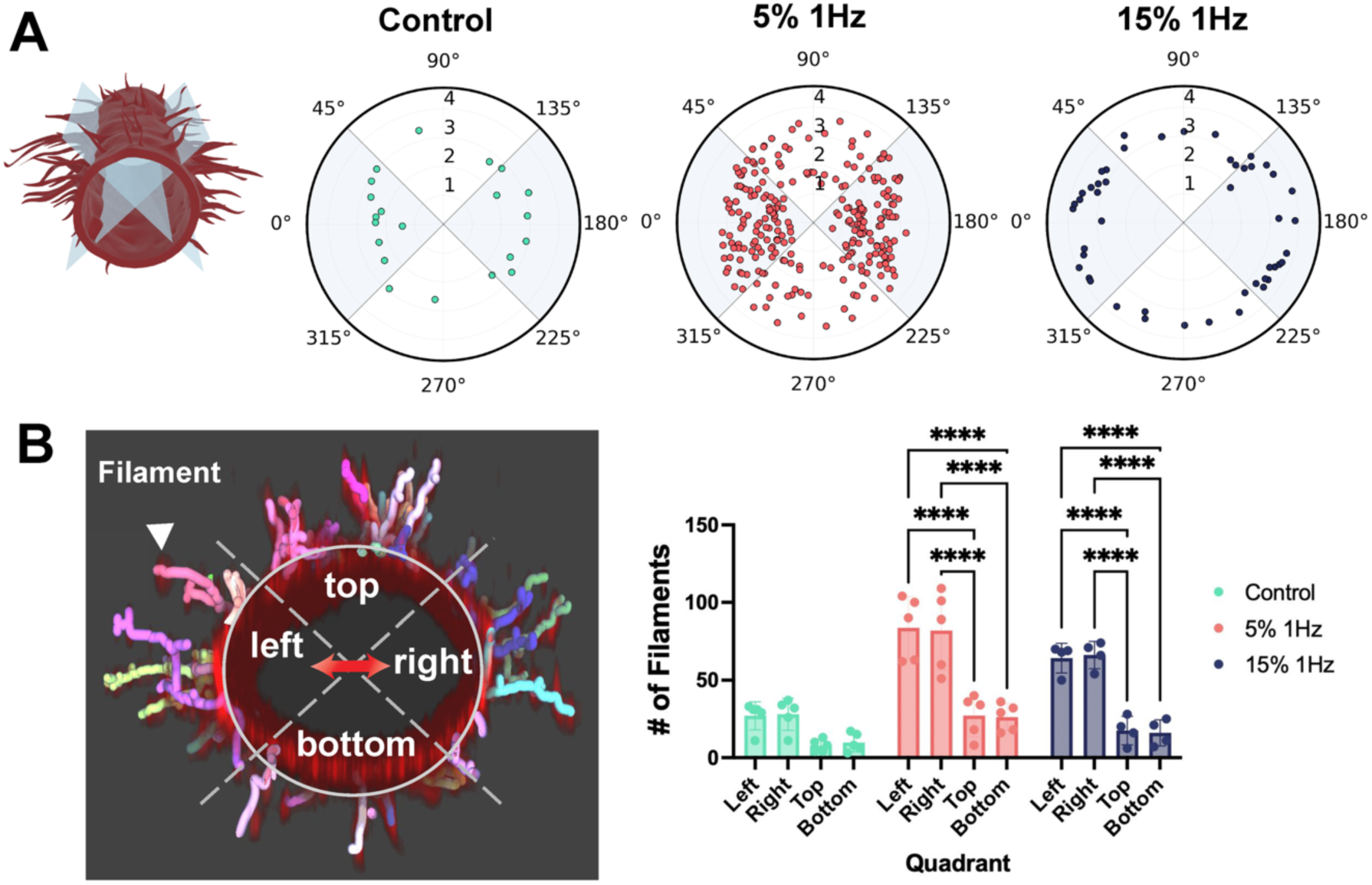
3D filament tracing and spatial orientation analysis of angiogenic sprouts. (*A*) Representative polar plots showing the orientation (θ) and length (log₁₀-transformed) of individual sprouts from one region of interest (ROI) under control, 5% 1 Hz, and 15% 1 Hz strain conditions. Each point corresponds to a single sprout. Shaded quadrants indicate regions aligned with the applied strain axis. (*B*) Spatial distribution of sprouts across vessel quadrants (left, right, top, bottom) for control, 5% dynamic, and 15% dynamic strain conditions. Quadrants were defined by dividing the vessel cross-section using angular boundaries at 45°, 135°, 225°, and 315°. Left: IMARIS 3D reconstruction of the vessel with individually color-labeled segmented filaments overlaid with the defined quadrant boundaries. The left-right axis corresponds to the direction of applied stimulation. Right: Quantification of the number of filaments within each vessel quadrant. (^ns^p > 0.05, *p < 0.05, **p < 0.01, ***p < 0.001, ****p < 0.0001, two-way ANOVA, n≥4 independent devices, one ROI per device).

### 4D spatiotemporal stimulation reprograms the direction of angiogenesis

We next explored whether dynamically reorienting the direction of mechanical stimulation over time could reprogram sprouting trajectories in 4D. We applied 15% strain at 1Hz along the x-axis for the first 3 days of stimulation, followed by reorientation of the strain direction to either the y- or z-axis for the subsequent 3 days. We refer to these conditions as XX stimulation (x-axis strain maintained throughout), XY stimulation (x- followed by y-axis strain), and XZ stimulation (x- followed by z-axis strain), respectively. To assess how these directional changes influenced sprouting behavior, we visualized sprouting morphology using 3D reconstructed z-stacks, capturing both top-down and cross-sectional views of the engineered vessels. Top-down images indicate that XY stimulation induced deflection of sprouts along the y-direction (green arrows), often forming L-shaped sprouts, in contrast to the primarily unidirectional sprouts observed under XX strain (Fig. 5A). Cross-sectional images indicate that XZ stimulation resulted in upward sprout extension along the z-axis (blue arrows), which was not observed in XX conditions (Fig. 5B).

**Figure 5.**
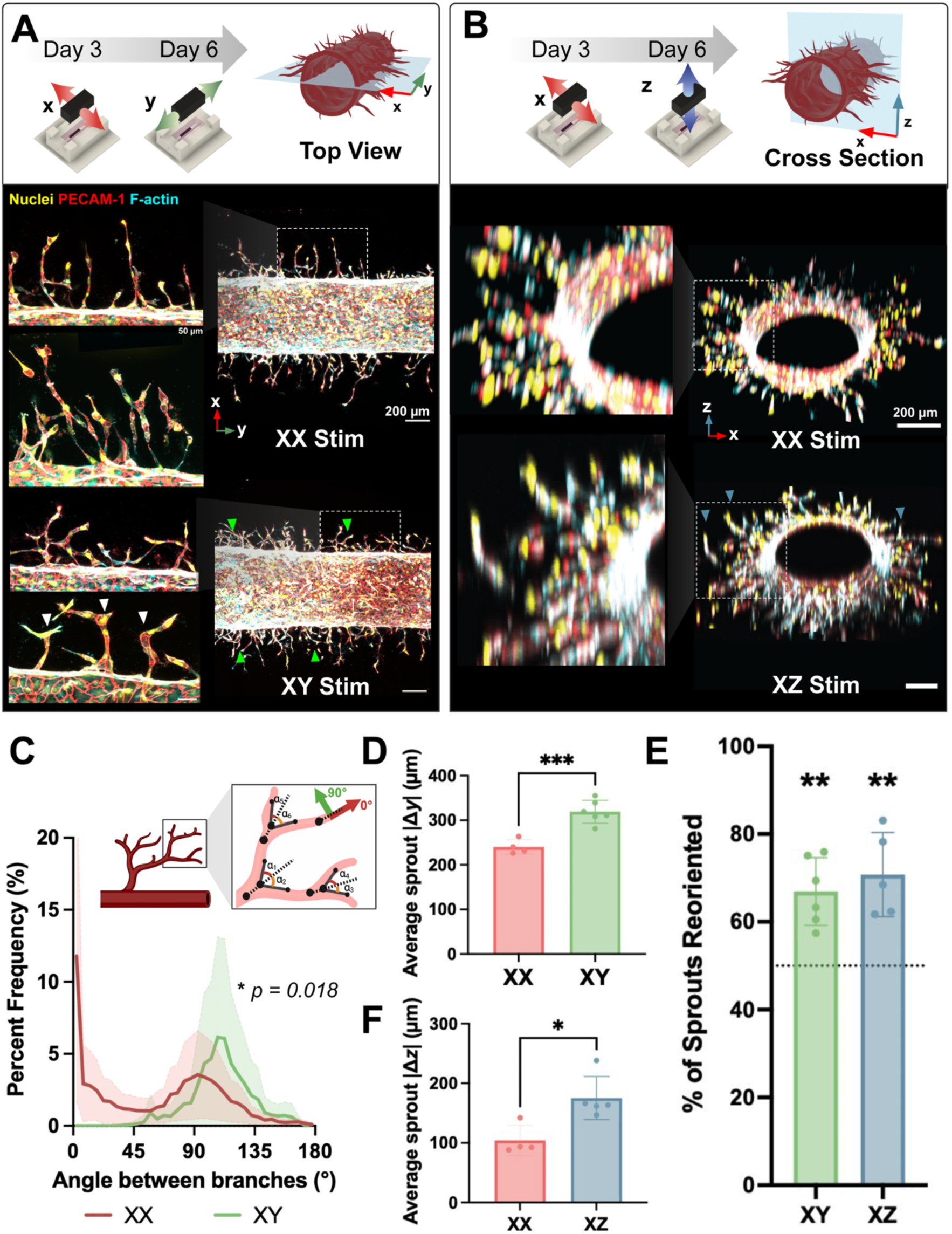
Dynamic reorientation of strain alters endothelial sprouting trajectory across spatial dimensions (*A*) Top-down 3D reconstructions of engineered vessels showing sprouting morphology under XX (x-axis strain maintained for 6 days) and XY (x-axis strain for 3 days followed by y-axis strain for 3 days) stimulation. Green arrows indicate deflection of sprouts along the y-direction under XY stimulation. Scale bars: 200 µm (large images), 50 µm (insets). (*B*) Cross-sectional 3D reconstructions of engineered vessels under XX and XZ (x-axis strain for 3 days followed by z-axis strain for 3 days) stimulation. Blue arrows indicate upward sprout extension along the z-axis under XZ stimulation. Scale bars, 200 µm (large images) (*C*) Distribution of branch angles measured between parent and daughter branches under XX and XY stimulation. The initial sprout from the parent vessel serves as the reference, and the angles of subsequent branches were analyzed. (p = 0.018, Kolmogorov–Smirnov test, n≥3 independent devices, one ROI per device). (*D*) Quantification of absolute change in y-position (|Δy|) per sprout under XX and XY stimulation. (^ns^p > 0.05, *p < 0.05, **p < 0.01, ***p < 0.001, Unpaired t-test, n≥4 independent devices, one ROI per device). (*E*) Percentage of sprouts classified as reoriented under XY and XZ stimulation relative to XX controls, defined using axis-specific displacement thresholds (|Δy| or |Δz| greater than the mean value under XX). Dashed line indicates 50%. (^ns^p > 0.05, *p < 0.05, **p < 0.01, n≥5 independent devices, one ROI per device) (*F*) Quantification of absolute change in z-position (|Δz|) per sprout under XX and XZ stimulation (^ns^p > 0.05, *p < 0.05, **p < 0.01, Unpaired t-test, n≥4 independent devices, one ROI per device).

We next examined whether L-shaped sprouts reflected additive extension along a newly imposed strain axis or remodeling of pre-existing growth trajectories. We longitudinally tracked changes in x- and y-displacement (Δx and Δy) of sprouts over the sequential stimulation period (Fig. S6A; Movie S7). During initial X stimulation (days 3–6), sprouts exhibited predominant increases in Δx with mild changes in Δy. Upon switching to Y stimulation (days 6–9), Δx plateaued without evidence of regression, while Δy increased. These results indicate that directional reprogramming reflects additive extension along the new strain axis rather than loss of prior x-directed growth.

We quantified how strain direction affects sprouting geometry by comparing XY and XZ conditions to XX stimulation. To assess XY stimulation, we quantified the angle between parent and daughter branches, using the initial sprout as the reference and measuring only subsequent branching events. Angles closer to 0° indicate growth along the x-axis, while angles near 90° represent perpendicular branching along the y-axis (Fig. 5C). Under XX stimulation, branch angles followed a bimodal distribution with a dominant peak at 2.5° and a secondary peak at 92.5°, consistent with predominantly x-direction sprouting. In contrast, XY produced a significantly different distribution of branch angles with a unimodal peak centered at 107.5°, indicating directional reprogramming of sprout trajectories in response to y-axis strain.

To quantify directional deflection under XY stimulation, we measured the absolute change in y-position (|Δy|) for each sprout and found a 1.3-fold increase in XY versus XX conditions (p = 0.0007), further confirming sprout deflection along the y-axis (Fig. 5D). Using the mean |Δy| under XX stimulation as a reference threshold, sprouts exhibiting |Δy| values greater than this baseline were classified as reoriented. Using this threshold, 67% of sprouts under XY stimulation were classified as reoriented, significantly greater than 50% (Fig. 5E, p<0.01). Among reoriented sprouts, left–right deflection was not significantly biased (Fig. S6B, 53.8% left vs. 46.2% right; p > 0.05), consistent with the bidirectional lateral strain imposed during XY stimulation.

To quantify the effects of XZ stimulation, we extracted the absolute change in z-position (|Δz|) for each sprout (Fig. 5F). Compared to XX stimulation, XZ stimulation resulted in a 1.7-fold increase in z-axis displacement. Using the mean |Δz| under XX as the reference threshold, 71% of sprouts were classified as reoriented under XZ stimulation, significantly exceeding 50% (Fig. 5E, p<0.01). In contrast to XY stimulation, vertical displacement under XZ stimulation was strongly biased in a single direction, with 98.7% of sprouts exhibiting positive z-deflection and only 0.3% deflecting downward (Fig. S4C, p<0.0001), consistent with the unidirectional upward strain imposed in this configuration.

In contrast to sprouting directionality, permeability measurements revealed no significant differences among the multi-directionally stimulated groups (XX, XY, XZ; p > 0.05) (Fig. S4D), indicating that vascular barrier function is independent of stimulation orientation in our platform. Consistent with the strain-induced permeability changes observed in Fig. 2, these results show that our 3D stimulation scheme can robustly reprogram sprout trajectories without affecting vessel integrity.

### RNA sequencing identifies transcriptional changes associated with mechanically stimulated angiogenesis

Given the distinct morphological and functional changes observed under mechanical stimulation, we investigated the molecular mechanisms underlying these strain-induced behaviors. Since cell-ECM interactions can strongly influence endothelial behavior, we first examined whether mechanical actuation altered the collagen hydrogel’s fibrillar architecture. Collagen fibers were visualized using the Rhobo6 fluorescent ECM-binding probe (35), and fiber orientation was quantified in the region between the magnetic actuator and the vessel. No significant differences in collagen fiber orientation were observed between unstimulated (0% strain) and dynamically actuated (15% 1Hz) conditions (Fig. S7A). These results indicated that dynamic magnetic matrix actuation did not significantly alter ECM microstructure, suggesting that observed endothelial responses could not be attributed to matrix remodeling.

We thus performed bulk RNA sequencing (RNA-seq) to compare gene expression between the unstimulated control, 5% 1Hz and 15% 1Hz strain conditions. Principal component analysis revealed clear separation between stimulated and control samples, while 5% 1Hz and 15% 1Hz strain conditions clustered closely together (Fig. 6A). Volcano plots for the pairwise comparisons highlighted up-regulated (red) and down-regulated (blue) differentially expressed genes (DEGs) (p. adjust < 0.05, |log_2_ fold change| > 0.7) (Fig. 6B; Fig. S8A). Differential expression analysis identified 570 DEGs under 15% 1Hz strain and 504 DEGs under 5% 1Hz strain, with 387 genes (56.4%) shared between both groups (Fig. S8B). Hierarchical clustering of Hallmark angiogenesis genes further showed that both strain conditions clustered together and were distinct from the unstimulated control, with upregulation of pro-angiogenic genes including *VEGFA, FGFR1, PTK2,* and *PDGFA* (Fig. 6C) (36).

**Figure 6.**
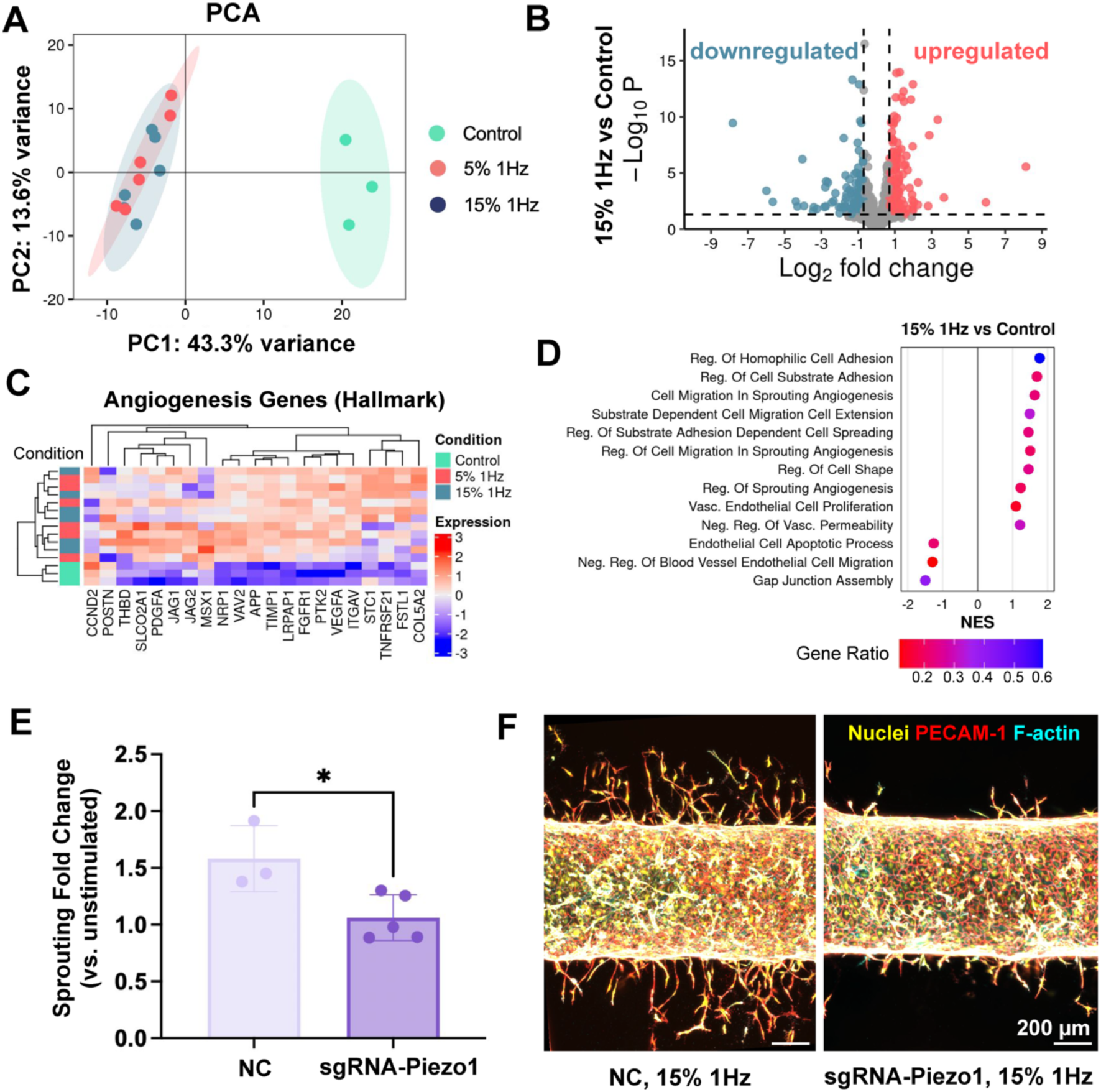
Strain-induced transcriptional reprogramming and functional validation (*A*) Principal component analysis (PCA) of bulk RNA-seq samples from unstimulated control, 5% 1Hz, and 15% 1Hz strain conditions. Strain groups (5% 1Hz and 15% 1Hz) had 5 samples with 2 pooled biological replicates per sample. Control group had 3 samples with 2 pooled biological replicates per sample. (*B*) Volcano plots showing upregulated (red) and downregulated (blue) differentially expressed genes (adjusted p < 0.05, |log₂ fold change| > 0.7) for 15% 1Hz relative to control. (*C*) Hierarchical clustering heatmap of Hallmark angiogenesis genes across unstimulated control, 5% 1Hz, and 15% 1Hz samples. (*D*) Gene set enrichment analysis (GSEA) of biological processes under 15% strain, ranked by normalized enriched score (NES). (*E)* Fold change in total sprouting under 15% 1 Hz stimulation relative to the unstimulated control for negative control (NC) and Piezo1 functional knockdown (sgRNA-Piezo1) conditions. (^ns^p > 0.05, *p < 0.05, Unpaired t-test, n≥3 independent devices, quantified across the entire device) *(F)* Representative confocal images of engineered vessels under 15% 1Hz stimulation in negative control (NC) and Piezo1 functional knockdown (sgRNA-Piezo1) conditions. Nuclei (yellow), PECAM-1 (red), and F-actin (cyan). Scale bar, 200 μm.

Gene set enrichment analysis (GSEA) was then used to interpret the functional significance of these transcriptional changes. Across both mechanically stimulated conditions, pathways associated with angiogenesis and cytoskeletal regulation exhibited positive normalized enrichment scores (NES > 1), including regulation of cell-substrate adhesion, cell migration in sprouting angiogenesis, endothelial cell proliferation, and regulation of cell shape, whereas pathways related to vascular permeability, apoptosis, and blood vessel regression showed negative enrichment (NES < 1) (Fig. 6D; Fig. S8C).

To further identify specific mechanosensitive mediators of this response, we explored a subset of genes with established roles in mechanosensing, transcriptional regulation, ECM remodeling, and barrier stabilization (Fig. S8D). Among the mechanosensors, the stretch-inducible adaptor *ANKRD1* was upregulated under 5% and 15% strain, consistent with its known roles in endothelial migration, survival, and neovascularization during tissue repair (37). The focal adhesion gene *ZYX* was strongly expressed under both 5% and 15% strain (p_adj_ < 0.001), and has been shown to transduce mechanical signals to the nucleus in endothelial cells and regulate stretch-sensitive gene expression (38, 39). Within sprouting-associated transcriptional regulators, *FOSB* was significantly upregulated in both 5% and 15% strain conditions (p_adj_ < 0.05 and < 0.01, respectively). As a mechanosensitive AP-1 family transcription factor, *FOSB* is upregulated by cyclic stretch in endothelial cells and contributes to AP-1 complexes that regulate endothelial genes involved in angiogenesis and vascular remodeling (39). The ECM and migration group showed robust gene upregulation across both strain magnitudes, with *ABI3BP*, *SPOCK1*, *COL8A1*, and *CTTN* all significantly upregulated (padj < 0.001) relative to control, suggesting enhanced substrate engagement and focal-adhesion remodeling that could facilitate sprouting behavior (40–43). In particular, *CTTN*, which localizes to adherens junctions and links actin dynamics to barrier regulation, may further contribute to strain-induced strengthening of endothelial junctions (42, 44). Supporting this, the upregulation of *SERPINE2* and downregulation of *P2RY1* align with our functional findings of strengthened barrier integrity under strain, consistent with transcriptional patterns that may support junctional stability while enabling controlled endothelial remodeling (45, 46). Together, these results may provide insight into a coordinated endothelial transcriptional program that integrates mechanosensing, angiogenic activation, and barrier stabilization under dynamic mechanical strain.

To assess the role of mechanosensitive signaling in strain-induced sprouting, we investigated the stretch-activated ion channel Piezo1. Piezo1 knockdown efficiency in HUVECs generated by Cas9–RNP nucleofection was confirmed by qRT-PCR 72 h post-transfection, demonstrating significant reduction in Piezo1 mRNA normalized to β-actin (Fig. S8E). We then compared total angiogenic sprouting under 15% 1Hz stimulation between negative control (NC) and Piezo1 knockdown (sgRNA-Piezo1) vessels.

In NC vessels, 15% stimulation induced a 1.6-fold increase in total sprouting relative to unstimulated controls, whereas Piezo1 knockdown vessels exhibited only a 1.1-fold increase (p < 0.05; Fig. 6E). Representative confocal images were consistent with these results, revealing decreased sprouting in sgRNA-Piezo1 vessels compared to NC vessels under 15% stimulation (Fig. 6F). These results demonstrate that Piezo1 loss significantly attenuates strain-induced sprouting. Characterizing Piezo1 effects on vessel barrier function, diffusive permeability measurements using 10 kDa dextran showed that 15% strain decreased permeability relative to unstimulated baseline in both NC and Piezo1 knockdown vessels (Fig. S8F). Together, these results suggest that Piezo1 regulates strain-induced sprouting but not strain-induced barrier stabilization.

## Discussion

Angiogenesis occurs in mechanically dynamic environments where tissue deformation, blood flow-induced stretch, and interstitial pressure continuously remodel the vasculature(15). While previous studies have identified mechanical forces as regulators of endothelial behavior(47), systematic investigation of how strain magnitude, frequency, and spatiotemporal patterning modulate 3D angiogenic sprouting and barrier function has been limited. Previous studies have shown that cyclic mechanical strain influences endothelial cell morphology and orientation (48). Yet how temporal dynamics, strain magnitude, and strain direction collectively influence directional angiogenic sprouting in 3D remains poorly understood.

Conventional global stretching platforms impose uniform deformation across an entire construct, restricting directional control of mechanical cues. In contrast, our magnetically actuated system enables programmable strain along defined spatial planes (x, y, and z), allowing axis-specific modulation of angiogenic patterning. In our study, strain magnitudes of 5–15% were chosen to match the physiological range of tensile loading experienced by blood vessels in native tissues.(16) Dynamic stimulation at 1Hz was employed to recreate the oscillatory mechanical environment characteristic of living tissue ECMs, critical for vascular remodeling.(49)

Our systematic analysis revealed that mechanical stimulation, regardless of strain magnitude (5% or 15%) or frequency (static or 1 Hz), reduced permeability to 10- and 40-kDa dextran tracers by approximately 2-fold and 1.5-fold, respectively. These results suggest that physiological strain engages a threshold-dependent mechanotransduction mechanism that enhances barrier integrity through actin cytoskeletal reorganization (50, 51). In contrast to 2D systems where uniaxial strain drives perpendicular alignment (52), endothelial cells in our 3D vessels experience tensile loading that promotes longitudinal junction organization. Notably, our endothelial junction orientation analysis was based on z-projections of the parent vessel, which may not fully capture organization at the vessel-gel interface where sprouting occurs, motivating future studies that examine strain-induced junctional remodeling at this interface.

In contrast to the magnitude-independent barrier response, sprouting morphology exhibited pronounced sensitivity to strain magnitude. While all mechanical stimulation increased total sprouting length, 5% dynamic strain maximized sprout number (density), whereas 15% dynamic strain produced the longest individual sprouts. This two-mode angiogenic response is consistent with previous work showing that moderate strain activates vessel formation pathways, whereas higher strain shifts the response toward MMP-driven matrix remodeling. (14) The shared upregulation of ECM remodeling genes (*ABI3BP, SPOCK1, COL8A1, CTTN*) across both 5% and 15% conditions indicates a common pro-angiogenic transcriptional program activated by dynamic mechanical stimulation. We note that the plateau in sprouting may also reflect matrix-dependent constraints in collagen, and testing more bioactive matrices such as fibrin or defined synthetic ECMs will be an important next step to examine longer-term lumen maturation under mechanical stimulation.(31, 32) Sprouts formed under stimulated conditions showed capillary-scale architectures, a size range that remains challenging to directly pattern using most 3D bioprinting strategies. This underscores the potential of dynamic mechanical stimulation as a complementary approach for organizing microvascular networks.

Leveraging quantitative mapping of sprouting trajectories in 3D, our results further demonstrate that mechanical strain provides a spatially instructive cue that biases angiogenic sprouting along the axis of deformation. Unstimulated vessels exhibited a slight but statistically insignificant baseline anisotropy, potentially due to subtle geometric influences of the rectangular device chamber and collagen casting architecture. When applied dynamically, unidirectional strain amplified this directional bias and induced a significant increase in sprouting along the strain axis, with 3.0-fold and 2.4-fold increases in sprout number for 5% and 15% conditions, respectively, relative to unstimulated controls. A key next step will be to systematically vary mechanical stimulation duration to define how timing shapes the initiation, progression, and maturation of strain-induced sprouts.

A unique contribution of our study is demonstrating how dynamically changing the axis of stimulation reprograms sprouting trajectories over time. Sequential application of strain along different axes, including y-and z-axis loading, produced L-shaped sprouts and deflection along the y- or z-direction, indicating that endothelial cells continuously adapt to evolving mechanical environments rather than adhering to a predetermined growth program. Notably, our findings demonstrate that directional reprogramming occurs in the majority of sprouts under dynamic strain reorientation, rather than representing a rare or isolated phenomenon. While previous studies have shown that the direction of static stretch or flow can influence sprouting orientation (53, 54), the ability to temporally redirect angiogenic sprouting has not been demonstrated in previous angiogenesis models, to our knowledge. The ability to steer 3D sprouting through controlled multi-directional strain, without measurable changes in permeability across x-, y-, and z-stimulated conditions, indicates continuous mechanosensing through focal adhesion components such as *ZYX* and dynamic actin remodeling mediated by *CTTN* (38, 42, 44).

To uncover the molecular basis of mechanically regulated angiogenesis, we performed transcriptomic profiling of endothelialized vessels subjected to dynamic strain. Bulk RNA sequencing revealed transcriptional remodeling in response to both 5% and 15% strain, with 387 shared DEGs (56.4% overlap). We observed minimal transcriptional divergence between 5% and 15% strain conditions, potentially because both magnitudes fall within a physiological range sufficient to activate a common mechanotransduction program. GSEA revealed robust enrichment of pathways associated with cell-substrate adhesion, cell migration in sprouting angiogenesis, endothelial cell proliferation, and regulation of cell shape, while pathways related to vascular permeability, apoptosis, and vessel regression were suppressed. This transcriptional signature aligns with our functional observations of enhanced sprouting and strengthened barrier function. Upregulation of VEGFA, FGFR1, PTK2 (FAK), and PDGFA in stimulated vessels is consistent with activation of canonical pro-angiogenic signaling pathways downstream of VEGF and FGF, in line with prior work showing that tensile forces regulate 3D vascular network morphogenesis through multiple autocrine and paracrine mechanisms. (18, 55)

To further explore candidate mechanosensors, we performed a functional knockdown of Piezo1, a well-established endothelial mechanosensor (56, 57). Although Piezo1 transcript levels were not significantly altered in our RNA-seq dataset, its contribution to strain sensing may be regulated by force-dependent channel gating or membrane tension rather than transcriptional upregulation (58, 59). Loss of Piezo1 reduced strain-induced sprouting, indicating that Piezo1 contributes to mechanotransduction pathways regulating angiogenesis. Interestingly, strain-dependent reductions in permeability were preserved in Piezo1-deficient vessels, suggesting that barrier stabilization under dynamic strain likely involves additional or parallel mechanosensitive pathways. Together, these findings support a model in which dynamic mechanical stimulation engages multiple mechanotransduction nodes, with Piezo1 representing one contributor among multiple mechanosensors. The modular design of our platform enables systematic genetic perturbation of candidate mechanosensors, providing a framework to dissect how force-sensing pathways regulate 3D vascular structure and function within a programmable mechanical environment.

Our study provides important insights into mechanically regulated angiogenesis; however, several limitations should be considered. Our human vessel-on-chip platform currently only includes endothelial cells, motivating future integration of smooth muscle cells, pericytes, and fibroblasts to better mimic the cellular complexity of native vasculature (60). In this work, we primarily focused on physiological strain magnitudes (5–15%) and a frequency of 1 Hz, relevant to oscillatory ECM environments of mechanically active tissues (e.g. heart, lungs, and muscle) (61). The response to pathological strains (>20%) or higher-frequency stimulation (>2 Hz) occurring in trauma or disease states thus remains unexplored (62, 63). Additionally, we conducted all experiments under static media culture conditions without incorporating physiological perfusion to isolate the effects of matrix-applied strain from confounding flow-derived forces such as shear stress. While flow contributes to cell–cell junction organization, permeability, and angiogenic gene expression (64), decoupling these variables allowed us to specifically examine the influence of strain on angiogenesis. Although cyclic deformation at 1Hz may generate transient flow with minimal, periodic shear within vessels, we observed no induction of shear-responsive genes (*KLF2, eNOS*) (52), whereas mechanical stimulation upregulated ECM/migration genes (*SPOCK1, COL8A1, CTTN*) and mechanosensitive markers (*ZYX, ANKRD1*), consistent with strain-driven rather than flow-driven signaling. We also acknowledge that in the absence of continuous flow, periodic media changes may have led to transient nutrient or oxygen gradients, which may influence the local microenvironment and contribute to depth-dependent sprouting behavior.

We observed that collagen fiber architecture remained unchanged across all mechanical conditions in our system, in contrast to other studies reporting mechanical remodeling of ECM fibers under strain (65). This is consistent with our prior work using magnetic matrix actuation platforms, where fibrin-based matrices also showed no reorganization under dynamic strain (27), suggesting that not all ECM materials undergo mechanically induced remodeling. Furthermore, given the lack of fibroblast-derived MMP activity in our endothelial-only model(66), the cellular and transcriptional changes observed cannot be attributed to matrix remodeling. While our RNA-seq revealed high transcriptional overlap between 5% and 15% strain conditions, the specific mechanisms by which strain magnitude differentially shapes sprouting phenotypes require further investigation and loss-of-function studies. Single-cell and spatial transcriptomic approaches may also reveal endothelial subpopulation heterogeneity in mechanical sensing and how these responses are spatially organized in 3D.

In conclusion, our findings demonstrate that mechanical forces regulate angiogenesis through coordinated effects of strain magnitude, frequency, and 4D spatiotemporal patterning. Dynamic strain at physiological magnitudes enhances endothelial alignment, sprouting, and barrier function through activation of mechanosensors, transcription factors, and ECM remodeling genes, with strain magnitude influencing whether angiogenic responses favor sprout initiation (5%) or elongation (15%). Despite substantial changes in vascular morphology, barrier function remained intact across all mechanical stimulation conditions, underscoring the potential of multi-axial actuation to engineer spatially organized yet functionally robust vascular networks. Our study is the first to demonstrate that sequential, multi-directional mechanical stimulation can reprogram angiogenic trajectories over time, enabling dynamic control of sprouting geometry across all three spatial dimensions. These results build upon previous work in 2D and uniaxial systems by extending the concept of strain-guided angiogenesis into a fully 3D microenvironment with programmable mechanical inputs.

Looking forward, our programmable 4D mechanical stimulation platform complements existing biochemical approaches for vascular patterning. Unlike diffusive growth factor gradients, which are challenging to control with precise spatiotemporal resolution in 3D tissues, defined strain fields provide a reconfigurable mechanical cue that can direct angiogenic growth in real-time. This capability opens opportunities to engineer hierarchical vascular networks with defined geometry and orientation that remain challenging to achieve using self-organization or static pre-patterning alone. Beyond tissue assembly for applications in regenerative medicine, our platform offers a controlled system to model mechanically dysregulated angiogenesis in disease, including tumor-associated stiffening, fibrosis-driven matrix contraction, and hypertensive vascular remodeling. Coupling programmable mechanical inputs with genetic or pharmacologic perturbations further enables systematic interrogation of how multiple mechanosensors and downstream signaling pathways coordinate to regulate vascular growth.

## Materials and Methods

Detailed materials and methods are available in SI Materials and Methods.

### Vessel-on-a-chip fabrication and cell seeding

The culture chamber was assembled from four polymethyl methacrylate (PMMA) side plates bonded with acrylic adhesive. A glass rod was inserted into the central channel, and the device was cast with 2.4 mg/mL collagen. A BSA-treated 3D-printed stamp was used to create a void for the magnetic actuator. Next, an additional collagen layer was cast. Following gelation, the rod was removed to form a hollow lumen. HUVECs were injected into the channel, and the device was rotated for 2 h to line endothelial cells uniformly. Vessel-on-a-chip devices were maintained in 6-well plates with 5 mL of medium, which was replaced every 2 days.

### High-throughput automated stimulation setup

A 3-axis CNC actuator system was adapted to provide programmable motion in X, Y, and Z directions. A 3D-printed holder positioned an array of neodymium magnets above vessel-on-a-chip devices within standard 6-well plates, enabling G-code–controlled magnetic actuation to apply tensile strain to the collagen matrix. The system accommodated up to three plates simultaneously and was operated at 37 °C.

### Permeability measurement

Permeability was assessed on day 7 via perfusion of fluorescent dextran. Dextran diffusion from the parent vessel into the ECM was imaged by confocal microscopy. The diffusive permeability coefficient (P_D_) was calculated from fluorescence intensity changes in the vessel and ECM using the mass-conservation relation J = P_D_ (c_vessel_ - c_ECM_), where J is the mass flux, c_vessel_ concentration in the parent vessel, and c_ECM_ is the concentration in the surrounding ECM.(67)

### Orientation analysis

Endothelial cell orientation was quantified from maximum-intensity projections of PECAM-1–stained vessels. Orientation distributions were analyzed in ImageJ using the *OrientationJ* plugin, which maps fiber directionality based on local image structure(68). Collagen fibers, stained with Rhobo-6(35), were analyzed using the same workflow to compare matrix alignment across strain conditions.

### RNA extraction, sequencing, and analysis

Cells were isolated by digesting gels with 0.5 mg/mL Liberase TM for 20 min at 37 °C, pooled from two devices, and RNA was extracted using the RNeasy Mini Kit. Libraries were prepared and sequenced at the MIT BioMicro Center, and RNAseq data were processed with the nf-core/rnaseq pipeline using Salmon for quantification and DESeq2 for differential expression.

### Generation of Piezo1 knockdown cells

Piezo1 knockdown HUVECs were generated by nucleofection of Cas9–sgRNA ribonucleoprotein complexes targeting Piezo1, followed by seeding into microfluidic devices for experiments. Knockdown efficiency was validated by qRT-PCR.

### Statistical analysis

Error bars represent mean ± standard deviation; statistical analyses were performed in GraphPad Prism 10 using two-tailed unpaired t-tests or one-/two-way ANOVA with Tukey’s multiple comparisons test (*p<0.05, **p<0.01, ***p<0.001, ****p<0.0001, ns p>0.05).

## Acknowledgments

This work was supported by a DoD Army Research Office Early Career Program and PECASE grant (W911NF-22-1-0126, awarded to R.R.) and a DoD DURIP Program grant (W911NF-24-1-0106, awarded to R.R.). S.K. acknowledges support from the Natural Sciences and Engineering Research Council of Canada (NSERC) Postdoctoral Fellowship, and J.S. acknowledges support from the National Science Foundation Graduate Research Fellowship Program (NSF-GRFP). We thank Dr. Kayvon Pedram for assistance with Rhobo6 imaging and Dr. Zhengpeng Wan for providing the cell line. We acknowledge the RNA sequencing support provided by the MIT BioMicro Center and imaging support from the Koch Institute Microscopy Core Facility. We also thank Sonika Kohli and Vesper Evereux for their assistance.

## References

1. N. Zhao, A. F. Pessell, N. Zhu, P. C. Searson, Tissue-Engineered Microvessels: A Review of Current Engineering Strategies and Applications. Adv. Healthc. Mater. 13, 2303419 (2024).

2. T. Rademakers, J. M. Horvath, C. A. van Blitterswijk, V. L. S. LaPointe, Oxygen and nutrient delivery in tissue engineering: Approaches to graft vascularization. J. Tissue Eng. Regen. Med. 13, 1815–1829 (2019).

3. R. H. Heisser, M. Bawa, J. Shah, A. Bu, R. Raman, Soft Biological Actuators for Meter-Scale Homeostatic Biohybrid Robots. Chem. Rev. 125, 3976–4007 (2025).

4. Y. Wang, M. Keshavarz, P. Barhouse, Q. Smith, Strategies for Regenerative Vascular Tissue Engineering. *Adv*. Biol. 7, 2200050 (2023).

5. S. Landau, et al., Bioengineering vascularization. Development 151, dev204455 (2024).

6. O. Aydin, et al., Principles for the design of multicellular engineered living systems. APL Bioeng. 6, 010903 (2022).

7. D. B. Kolesky, K. A. Homan, M. A. Skylar-Scott, J. A. Lewis, Three-dimensional bioprinting of thick vascularized tissues. Proc. Natl. Acad. Sci. 113, 3179–84 (2016).

8. P. Carmeliet, R. K. Jain, Molecular mechanisms and clinical applications of angiogenesis. Nature 473, 298–307 (2011).

9. N. Walji, S. Kheiri, E. W. K. Young, Angiogenic Sprouting Dynamics Mediated by Endothelial-Fibroblast Interactions in Microfluidic Systems. *Adv*. Biol. 5, 2101080 (2021).

10. P. Chai, et al., GlycoRNA complexed with heparan sulfate regulates VEGF-A signalling. Nature 1–11 (2026). 10.1038/s41586-025-10052-8.

11. D. Rana, et al., Bioprinting of Aptamer-Based Programmable Bioinks to Modulate Multiscale Microvascular Morphogenesis in 4D. Adv. Healthc. Mater. 14, 2402302 (2025).

12. F. E. Freeman, et al., 3D bioprinting spatiotemporally defined patterns of growth factors to tightly control tissue regeneration. Sci. Adv. 6, eabb5093 (2020).

13. J. Flournoy, S. Ashkanani, Y. Chen, Mechanical regulation of signal transduction in angiogenesis. Front. Cell Dev. Biol. 10 (2022).

14. N. F. Jufri, A. Mohamedali, A. Avolio, M. S. Baker, Mechanical stretch: physiological and pathological implications for human vascular endothelial cells. Vasc. Cell 7, 8 (2015).

15. X. R. Lim, O. F. Harraz, Mechanosensing by Vascular Endothelium. Annu. Rev. Physiol. 86, 71–97 (2024).

16. S. Barrasa-Ramos, C. A. Dessalles, M. Hautefeuille, A. I. Barakat, Mechanical regulation of the early stages of angiogenesis. J. R. Soc. Interface 19, 20220360 (2022).

17. T. Korff, H. G. Augustin, Tensional forces in fibrillar extracellular matrices control directional capillary sprouting. J. Cell Sci. 112, 3249–3258 (1999).

18. D. Rosenfeld, et al., Morphogenesis of 3D vascular networks is regulated by tensile forces. Proc. Natl. Acad. Sci. 113, 3215–3220 (2016).

19. J. H.-C. Wang, P. Goldschmidt-Clermont, J. Wille, F. C.-P. Yin, Specificity of endothelial cell reorientation in response to cyclic mechanical stretching. J. Biomech. 34, 1563–1572 (2001).

20. R. Kaunas, S. Usami, S. Chien, Regulation of stretch-induced JNK activation by stress fiber orientation. Cell. Signal. 18, 1924–1931 (2006).

21. D. Ferrari, et al., Effects of biomechanical and biochemical stimuli on angio- and vasculogenesis in a complex microvasculature-on-chip. iScience 26, 106198 (2023).

22. A. Shimizu, et al., ECM-based microchannel for culturing in vitro vascular tissues with simultaneous perfusion and stretch. Lab. Chip 20, 1917–1927 (2020).

23. L. Debbi, et al., The Effect of Mechanical Loading on Sprouting Angiogenesis from Engineered Macro-vessel Model. Small Methods n/a, e00850 (2025).

24. M. A. Moreno-Mateos, et al., Magneto-mechanical system to reproduce and quantify complex strain patterns in biological materials. *Appl*. Mater. Today 27, 101437 (2022).

25. D. Garcia-Gonzalez, R. Raman, S. Schuerle, A. Tay, Magnetic Actuation for Mechanomedicine. Adv. Intell. Syst. 7, 2400638 (2025).

26. B. Rios, et al., Mechanically programming anisotropy in engineered muscle with actuating extracellular matrices. Device 1 (2023).

27. A. Bu, et al., Actuating Extracellular Matrices Decouple the Mechanical and Biochemical Effects of Muscle Contraction on Motor Neurons. Adv. Healthc. Mater. 14, 2403712 (2025).

28. D. J. Green, M. T. E. Hopman, J. Padilla, M. H. Laughlin, D. H. J. Thijssen, Vascular Adaptation to Exercise in Humans: Role of Hemodynamic Stimuli. Physiol. Rev. 97, 495–528 (2017).

29. K. J. Peyton, X. Liu, W. Durante, Prolonged Cyclic Strain Inhibits Human Endothelial Cell Growth. Front. Biosci. Elite Ed. 8, 205–212 (2016).

30. C. O’Connor, E. Brady, Y. Zheng, E. Moore, K. R. Stevens, Engineering the multiscale complexity of vascular networks. Nat. Rev. Mater. 7, 702–716 (2022).

31. J. Pauty, et al., A Vascular Endothelial Growth Factor-Dependent Sprouting Angiogenesis Assay Based on an In Vitro Human Blood Vessel Model for the Study of Anti-Angiogenic Drugs. eBioMedicine 27, 225–236 (2018).

32. J. W. Song, L. L. Munn, Fluid forces control endothelial sprouting. Proc. Natl. Acad. Sci. 108, 15342–15347 (2011).

33. V. van Duinen, et al., Perfused 3D angiogenic sprouting in a high-throughput in vitro platform. Angiogenesis 22, 157–165 (2019).

34. S. Zhu, et al., Magnetically Controlled Strategies for Enhanced Tissue Vascularization. Adv. Funct. Mater. 34, 2401856 (2024).

35. A. Fiore, et al., Live imaging of the extracellular matrix with a glycan-binding fluorophore. Nat. Methods 1–11 (2025). 10.1038/s41592-024-02590-2.

36. A. Liberzon, et al., The Molecular Signatures Database Hallmark Gene Set Collection. Cell Syst. 1, 417–425 (2015).

37. X. Xu, et al., Research progress of ankyrin repeat domain 1 protein: an updated review. Cell. Mol. Biol. Lett. 29, 131 (2024).

38. A. Wójtowicz, et al., Zyxin Mediation of Stretch-Induced Gene Expression in Human Endothelial Cells. Circ. Res. 107, 898–902 (2010).

39. Y. Fang, D. Wu, K. G. Birukov, Mechanosensing and Mechanoregulation of Endothelial Cell Functions. Compr. Physiol. 9, 873–904 (2019).

40. Q. Li, et al., COL8A1 Regulates Endothelial Phenotype in Inflammatory Endothelial-to-Mesenchymal Transition. Am. J. Physiol.-Heart Circ. Physiol. (2025). 10.1152/ajpheart.00339.2025.

41. L. Váncza, P. Tátrai, A. Reszegi, K. Baghy, I. Kovalszky, SPOCK1 with unexpected function. The start of a new career. Am. J. Physiol.-Cell Physiol. 322, C688–C693 (2022).

42. S. Moztarzadeh, et al., Cortactin is in a complex with VE-cadherin and is required for endothelial adherens junction stability through Rap1/Rac1 activation. Sci. Rep. 14, 1218 (2024).

43. D. A. Delfín, J. L. DeAguero, E. N. McKown, The Extracellular Matrix Protein ABI3BP in Cardiovascular Health and Disease. Front. Cardiovasc. Med. 6 (2019).

44. X. Sun, et al., Genetic and epigenetic regulation of cortactin (CTTN) by inflammatory factors and mechanical stress in human lung endothelial cells. Biosci. Rep. 44, BSR20231934 (2024).

45. L. Idir, et al., SerpinE2 deficiency exacerbates glomerular injury in diabetic nephropathy through dysregulated angiogenesis and inflammatory responses. Am. J. Physiol.-Ren. Physiol. 329, F548–F556 (2025).

46. J. K. Hennigs, et al., The P2-receptor-mediated Ca2+ signalosome of the human pulmonary endothelium - implications for pulmonary arterial hypertension. Purinergic Signal. 15, 299–311 (2019).

47. A. N. Ivanov, Yu. R. Chabbarov, Mechanisms of Physiological Angiogenesis. J. Evol. Biochem. Physiol. 59, 914–929 (2023).

48. I. S. Joung, M. N. Iwamoto, Y.-T. Shiu, C. T. Quam, Cyclic strain modulates tubulogenesis of endothelial cells in a 3D tissue culture model. Microvasc. Res. 71, 1–11 (2006).

49. P. M. Cummins, et al., Cyclic strain-mediated matrix metalloproteinase regulation within the vascular endothelium: a force to be reckoned with. Am. J. Physiol.-Heart Circ. Physiol. 292, H28–H42 (2007).

50. P. Keshavanarayana, F. Spill, A mechanical modeling framework to study endothelial permeability. Biophys. J. 123, 334–348 (2024).

51. A. Bancaud, et al., Intraluminal pressure triggers a rapid and persistent reinforcement of endothelial barriers. Lab. Chip 25, 2061–2072 (2025).

52. S. Chien, Mechanotransduction and endothelial cell homeostasis: the wisdom of the cell. Am. J. Physiol.-Heart Circ. Physiol. 292, H1209–H1224 (2007).

53. S. Das, A. Ippolito, P. McGarry, V. S. Deshpande, Cell reorientation on a cyclically strained substrate. PNAS Nexus 1, pgac199 (2022).

54. P. A. Galie, et al., Fluid shear stress threshold regulates angiogenic sprouting. Proc. Natl. Acad. Sci. 111, 7968–7973 (2014).

55. Y. C. Yung, J. Chae, M. J. Buehler, C. P. Hunter, D. J. Mooney, Cyclic tensile strain triggers a sequence of autocrine and paracrine signaling to regulate angiogenic sprouting in human vascular cells. Proc. Natl. Acad. Sci. 106, 15279–15284 (2009).

56. S. S. Ranade, et al., Piezo1, a mechanically activated ion channel, is required for vascular development in mice. Proc. Natl. Acad. Sci. 111, 10347–10352 (2014).

57. J. Li, et al., Piezo1 integration of vascular architecture with physiological force. Nature 515, 279–282 (2014).

58. C. D. Cox, et al., Removal of the mechanoprotective influence of the cytoskeleton reveals PIEZO1 is gated by bilayer tension. Nat. Commun. 7, 10366 (2016).

59. A. H. Lewis, J. Grandl, Mechanical sensitivity of Piezo1 ion channels can be tuned by cellular membrane tension. eLife 4, e12088 (2015).

60. H. Kim, T. Osaki, R. D. Kamm, H. H. Asada, Tri-culture of spatially organizing human skeletal muscle cells, endothelial cells, and fibroblasts enhances contractile force and vascular perfusion of skeletal muscle tissues. FASEB J. 36, e22453 (2022).

61. C. A. Dessalles, C. Leclech, A. Castagnino, A. I. Barakat, Integration of substrate- and flow-derived stresses in endothelial cell mechanobiology. *Commun*. Biol. 4, 1–15 (2021).

62. Y. Song, et al., The Molecular Mechanism of Aerobic Exercise Improving Vascular Remodeling in Hypertension. Front. Physiol. 13 (2022).

63. I. A. Tamargo, K. I. Baek, Y. Kim, C. Park, H. Jo, Flow-induced reprogramming of endothelial cells in atherosclerosis. Nat. Rev. Cardiol. 20, 738–753 (2023).

64. M. Cherubini, et al., Flow in fetoplacental-like microvessels in vitro enhances perfusion, barrier function, and matrix stability. Sci. Adv. 9, eadj8540 (2023).

65. M. G. McCoy, et al., Collagen Fiber Orientation Regulates 3D Vascular Network Formation and Alignment. ACS Biomater. Sci. Eng. 4, 2967–2976 (2018).

66. I. Stamenkovic, Extracellular matrix remodelling: the role of matrix metalloproteinases. J. Pathol. 200, 448–464 (2003).

67. W. J. Polacheck, M. L. Kutys, J. B. Tefft, C. S. Chen, Microfabricated blood vessels for modeling the vascular transport barrier. Nat. Protoc. 14, 1425–1454 (2019).

68. Z. Püspöki, M. Storath, D. Sage, M. Unser, “Transforms and Operators for Directional Bioimage Analysis: A Survey” in Focus on Bio-Image Informatics, W. H. De Vos, S. Munck, J.-P. Timmermans, Eds. (Springer International Publishing, 2016), pp. 69–93.

