## Supplemental Information for "4D force patterning enables spatial control of angiogenesis"

**This PDF file includes:**

Materials and Methods
Figures S1 to S8
Legends for Movies S1 to S7

**Other supporting materials for this manuscript include the following:**

Movies S1 to S7

### Materials and Methods

#### Device fabrication

The culture chamber was assembled from four polymethyl methacrylate (PMMA) side plates (⅛ inch thickness, McMaster Carr) using an acrylic adhesive (Weld-On 4). The two end wall plates have a 650μm diameter hole. All PMMA pieces were cut using a CO2 laser (Laser Cutter Epilog Fusion 40). The culture chamber was then bonded to a square glass coverslip (No. 2 glass, VWR) using double-sided adhesive tape (9474LE, 3M)(79). The internal length, width, and depth were 10mm, 4mm, and 3.175mm, respectively (Fig. S1A-B). The devices were then autoclaved for sterilization. Magnetic actuators were cut from commercial neodymium magnet sheets (thickness: 16 mil, Bandle B) using an electronic cutting plotter (Silhouette Cameo 4, Silhouette America).

#### Cell culture and seeding in the microfluidic device

ImHUVECs expressing blue fluorescent protein (BFP, passages 12–20) were cultured in VascuLife VEGF-Mv Endothelial medium (Lifeline Cell Technology), with medium changes every 2 days. Cells were detached with TrypLE Express (Gibco) upon reaching ~80% confluency and seeded into microfluidic devices.

The fabrication process of the vascular construct is shown in Figure S1. Briefly, a single glass rod (DWK Life Sciences) was inserted into the central channel of the device. Next, 180 μL of collagen solution (2.4 mg/mL, porcine tendon, Cellmatrix Type I-A) was prepared on ice and cast into the device. A BSA-treated 3D-printed stamp was then aligned with the device using the built-in features to create a void space for the magnetic actuator. The device was incubated for 1h at 37 ºC for collagen gelation. Next, the stamp was carefully removed from the device, and the magnetic actuator was positioned in the patterned void space, followed by the addition of 80 μL premixed collagen to sandwich the actuator. This top collagen layer was gelled at 37 °C for 1 h. The glass rod was then removed, and any residual liquid in the channel was aspirated with a pipette to yield a hollow lumen. A 10 μL suspension of ImHUVECs ($8 \times 10^6$ cells/mL) was injected into the lumen, and devices were incubated at 37 °C for 2 h on a Tube Revolver Rotator platform (Thermo Scientific) to promote uniform 3D lining of the channel. Each device was maintained in a 6-well plate (VWR) with 5 mL of culture medium to fully submerge the construct, and the medium was replenished every 2 days.

#### Strain characterization and high-throughput mechanical stimulation

Green fluorescent polystyrene microspheres (Fisher Scientific) were embedded within the hydrogel to track matrix deformation. Videos were recorded using an EVOS fluorescence microscope and processed using the authors' previously developed open-source computational framework (https://github.com/HibaKob/Raman_Manuscript_2023)(32) to calculate the subdomain-averaged strain in each row over time.

#### High-throughput automated stimulation setup

A custom-built high-throughput stimulation platform was developed by repurposing a benchtop three-axis linear actuator system originally designed for computer numerical control (CNC) milling (3020-PRO MAX V2, Genmitsu). The base assembly included motorized X, Y, and Z linear actuators that provided programmable motion in all three dimensions (motion range: X 300mm; Y 205 mm; Z 78 mm; positional accuracy ±0.1 mm). To adapt the system for magnetic stimulation experiments, a custom 3D-printed holder was designed and mounted onto the actuator arm to secure an array of up to nine external neodymium magnets (3896.66 Surface Gauss, 0.750" L x 0.375" W x 0.375" H, Radial Magnets) above the vessel-on-a-chip devices. A 3D-printed well plate insert was fabricated to position the chips precisely within standard 6-well plates. A multi-well plate holder was also added to the stage to accommodate up to three well plates simultaneously, enabling parallel mechanical stimulation experiments. The entire system was controlled via G-code instructions, providing programmable control over magnet motion parameters, including direction, distance, and frequency of actuation. During stimulation, magnets were displaced from a central neutral position above each device to actuate the embedded magnetic rod and introduce tensile strain into the surrounding collagen matrix, and were then returned to the neutral position to complete each cycle. Magnet motion was constrained to remain within this neutral-to-actuated range and did not extend past the midpoint position, resulting in a tension–relaxation cycle rather than alternating tension–compression. The automated setup was housed within a standard laboratory oven maintained at 37°C to ensure physiological conditions throughout the stimulation period.

**Permeability measurement**
To assess vascular barrier permeability, devices were perfused with fluorescent dextran (10 kDa Texas Red and 40 kDa Alexa Fluor 647, Invitrogen) on day 7. Permeability measurements were performed specifically on straight segments of the parent vessel, with regions of interest (ROIs) defined based on dextran fluorescence within the vessel wall, which did not include angiogenic sprout regions. Dextran diffusion from the parent vessel into the ECM was imaged in real-time using the Nikon AXR confocal microscope. Confocal z-stacks were acquired every 2 minutes over 10 minutes (10 kDa) and 30 minutes (40 kDa), and maximum intensity projection images were used for analysis. The diffusive permeability coefficient ($P_D$) was calculated in MATALB by analyzing the changes in fluorescence intensity in the vessel and ECM. The diffusion profile was fitted to a dynamic mass-conservation equation $J = P_D (c_{vessel} - c_{ECM})$, where J is the mass flux of dextran, $c_{vessel}$ dextran concentration in the parent vessel, and $c_{ECM}$ is the dextran concentration in the surrounding ECM, following established methods(41, 42).

**Immunofluorescence staining and microscopy**
To visualize endothelial sprouting, the cells in the devices were fixed with 4% paraformaldehyde for 20 minutes at room temperature, permeabilized with 0.2% Triton X-100/PBS for 5 minutes, and blocked in 1% BSA in 0.2% Triton X-100/PBS for 30 minutes. The cells were then washed three times with 0.2% Triton X-100/PBS for 5 minutes each and incubated overnight at 4°C with Alexa Fluor 488-conjugated mouse anti-human platelet/endothelial cell adhesion molecule-1 (PECAM-1, 1:300 dilution, Cell Signaling Technology). After overnight incubation, the cells were rinsed three times with 0.2% Triton X-100/PBS and incubated with Alexa Fluor Plus 555 Phalloidin (1:400 dilution, Thermo Fisher Scientific) for 20 minutes at room temperature. The cells were then washed three times with PBS and stored in PBS for immunofluorescence imaging. Fluorescence images were captured on a Nikon AXR point scanning confocal microscope using 4x, 10x, and 20x objectives (Nikon) and processed using the Nikon Denoise.ai software and ImageJ.

**Sprouting quantification**
Sprouting was quantified from confocal images of cells immunostained for F-actin. For each device, z-stacks spanning the full vessel length were acquired and reconstructed as maximum-intensity projections of the entire vessel. Quantification was performed on the full-vessel projection for each device (one full-vessel analysis per independent device is used for statistical analysis). Full vessel projections were thresholded to isolate vascular structures, despeckled to remove noise, and skeletonized. The skeletonized images were then analyzed using a custom graph-based algorithm implemented in Wolfram Mathematica. Each binary skeleton was converted to a morphological graph, where continuous vessel and sprout segments were represented as edges, and branch junctions (vertices with degree >2) and terminal sprout tips (vertices with degree=1) were represented as vertices.

The graph was decomposed into connected components relative to primary branch points, corresponding to the origins of sprouts from the parent vessel. For each connected component, sprouting metrics were calculated by selecting the parent branch point and computing Euclidean distances to all associated endpoints. The maximum sprout length was defined as the longest Euclidean distance from the parent vessel to a terminal endpoint, and the total sprout length as the sum of all distances within the component. All sprouting metrics were calculated from full-vessel reconstructions for each device.

To analyze daily changes in sprouting, images of the BFP-tagged nuclei signal across the full vessel were acquired. Maximum projection images were processed in Fiji using the Huang thresholding method to generate binary vessel masks. Since the BFP signal does not form a continuous structure suitable for skeletonization, the perimeters of the masks were extracted using the Analyze Particles function to determine the vessel boundary contour length, which served as a global metric of sprouting across the full vessel. For each device, contour lengths from each day were normalized to the value measured on the day stimulation began.

**Orientation analysis**
The orientation of endothelial cells within the vascular constructs was quantified using immunostaining for PECAM-1, a marker of cell–cell junctions. The entire constructs were imaged using a Nikon AXR confocal

microscope, and Z-stack images were acquired across the full height of the vessels. Maximum intensity projections of the Z-stacks were generated for analysis. ImageJ (NIH) with the OrientationJ plugin was used to measure the angular distribution of endothelial cells under different experimental conditions. (80) The resulting images were displayed in HSB color, where hue indicates orientation, saturation represents coherency, and brightness corresponds to the original signal intensity.

Collagen fibers were stained with Rhobo-6 (43) and imaged using confocal imaging. Fiber orientation was quantified using an identical OrientationJ workflow, and differences in alignment were compared across the various mechanical stimulation conditions.

**Image analysis and 3D vessel reconstruction**
Confocal microscopy images of immunofluorescently labeled vascular networks were acquired using the Nikon AXR confocal microscope. Image stacks were processed using Imaris image analysis software (Oxford Instruments, version 10.1). 3D reconstructions of the vascular structures were generated from the complete z-stacks to capture the full morphology and spatial organization of sprouting angiogenesis within the tissue constructs. The coordinates of all filament skeleton nodes were extracted using the "Filament Point Tracking" module, a MATLAB-integrated feature within Imaris, which exports the complete set of (x, y, z) coordinates for every segment. These coordinates were then used for downstream spatial analyses, including quantification of sprout positions and their growth directions. The Angle Measurement tool within Imaris was also employed to compute the angles between sprouts at branch points, providing quantitative information on the three-dimensional organization and branching orientation of sprouts. All quantitative data generated using the Imaris Filament Tracer, Filament Point Tracking, and Angle Measurement modules were exported as CSV files and subsequently processed and analyzed using MATLAB (MathWorks) and GraphPad Prism 10 to perform statistical analysis and generate graphical representations.

**RNA extraction, sequencing, and analysis**
Cells were extracted by removing the gel from the devices and digesting each gel in 500 µL of 0.5 mg/mL Liberase TM (Sigma Aldrich) in VascuLife Basal Medium for 20 minutes at 37°C on a revolver(81). Cells were pooled from 2 devices. RNA extraction was performed on the isolated cells using a RNeasy Mini Kit (Qiagen) following the manufacturer instructions. Bulk RNA sequencing was performed by the MIT BioMicro Center. Sequencing libraries were generated using the Low Input hs|mm ZapR protocol with rRNA depletion and strand-specific preparation, followed by quality assessment. Libraries were sequenced on a Singular G4 platform with 50 bp paired-end reads. Raw and processed RNA sequencing data have been deposited in the Gene Expression Omnibus (GEO) under accession number GSE310169.

Raw FASTQ files were processed using the nf-core/rnaseq pipeline (v3.17.0). Transcript abundances were quantified using Salmon with the GENCODE v47 human transcriptome as reference. Differential expression analysis was performed in DESeq2 (v1.42.1) with log2 fold-change shrinkage via the apeglm method, and statistical significance was determined using the Benjamini–Hochberg false discovery rate (FDR ≤ 0.05). Functional enrichment was performed with fgsea using MSigDB Hallmark and C5 Gene Ontology Biological process collections, and results were visualized in R (v4.3) using ggplot2 and ComplexHeatmap.

**Generation of Piezo1 knockdown cells**
Cas9-RNP complexes were used for generating Piezo1 knockdown (sgRNA-Piezo1) HUVECs. For $1\times10^6$ HUVECs, 20 pmol SpCas9 (Synthego, R20SPCAS9-MED) and 30 pmol Piezo1 sgRNA (AGGACCGCGCCGAGCACGTG, ordered from Synthego) were used. Nucleofection was performed in Amaxa Biosystems Nucleofector II (Program HUVEC). Immediately following nucleofections, cells were transferred to pre-warmed medium and seeded into microfluidic devices as described above for subsequent experiments.

Quantitative reverse transcriptase PCR (qRT-PCR) was performed to validate the gene editing efficiency using Brilliant II SYBR qRT-PCR one-step Master kit (600825, Agilent) on a CFX Connect Real-Time System (Bio-Rad). RNAs were extracted from HUVECs 72-hour post nucleofection. 60 ng of RNA was used in the reaction with 1.25 µM primers (F Piezo1: ACTTTGCCCTGTCCGCCT; R Piezo1:

GAAGAAGCCCCGCACAAAC; F actin: CACCATTGGCAATGAGCGGTTC; R β-actin: AGGTCTTTGCGGATGTCCACGT). The data were analyzed using the ΔΔCt method.

Total sprouting for Piezo1 knockdown and negative control vessels was quantified from confocal maximum-intensity projections of F-actin–stained full vessels, as described above. For each independent device, the entire vessel length was analyzed to extract total sprout length using the graph-based skeletonization workflow. For fold-change comparisons under 15% 1 Hz stimulation, total sprouting values were normalized to the mean total sprouting measured in the corresponding unstimulated control group. Statistical analysis was performed using one full-vessel measurement per independent device.

**Statistical analysis**
All error bars are shown as mean ± standard deviation, and sample sizes are reported in figure legends. Statistical analysis was performed using GraphPad Prism 10. Ordinary one-way or two-way ANOVA with Tukey's multiple comparisons test was applied to analyses involving more than two groups, and two-tailed unpaired t-tests were applied to analyses involving two groups. * indicates $p<0.05$, ** indicates $p<0.01$, *** indicates $p<0.001$, **** indicates $p<0.0001$, and ns indicates $p>0.05$.

**Figures**

A

C

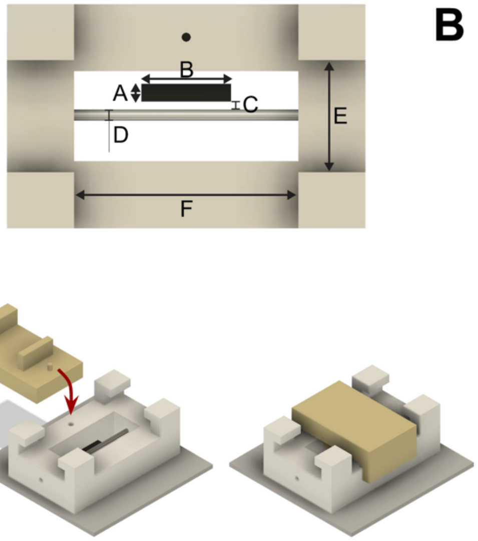


B

| Label | Description | Length (mm) |
|---|---|---|
| A | Microactuator width | 0.8 |
| B | Microactuator length | 4 |
| C | Actuator to vessel distance | 0.5325 |
| D | Vessel diameter | 0.635 |
| E | Device inner width | 4 |
| F | Device inner length | 10 |
| – | Device inner depth | 3.175 |

D

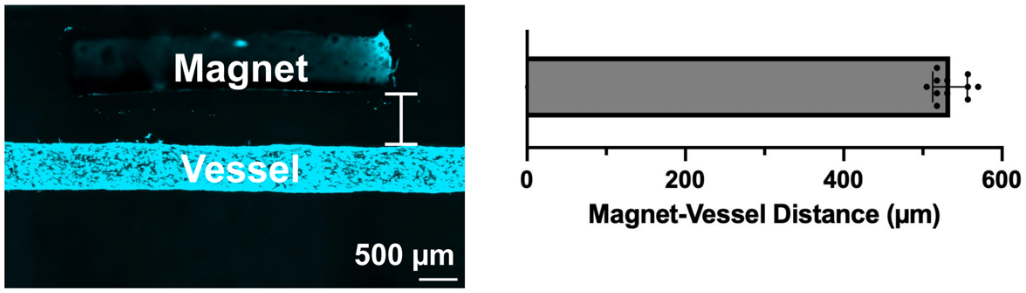


**Fig. S1.** Device geometry and local strain mapping of the magnetically actuated vessel-on-a-chip platform. (*A*) Top-down schematic of the vessel-on-a-chip device showing key geometric parameters. Lettered labels correspond to dimensions listed in panel B, indicating device geometry, actuator size, and actuator-to-vessel spacing. (*B*) Table summarizing geometric parameters and dimensions of the vessel-on-chip device. (*C*) Schematic of the custom 3D-printed stamp used to define a dedicated actuator cavity within the collagen matrix. The stamp contains an alignment pin that interfaces with a corresponding slot in the acrylic device (red arrow) to ensure reproducible positioning of the magnetic actuator relative to the engineered vessel. (*D*) Maximum-intensity projection of a confocal z-stack showing the magnetic actuator positioned parallel to the engineered vessel within the collagen matrix. Quantification of actuator–vessel spacing across independent devices is shown below (n = 11 independent devices). Mean edge-to-edge distance = 535 ± 20 µm. Scale bar is 500 µm.

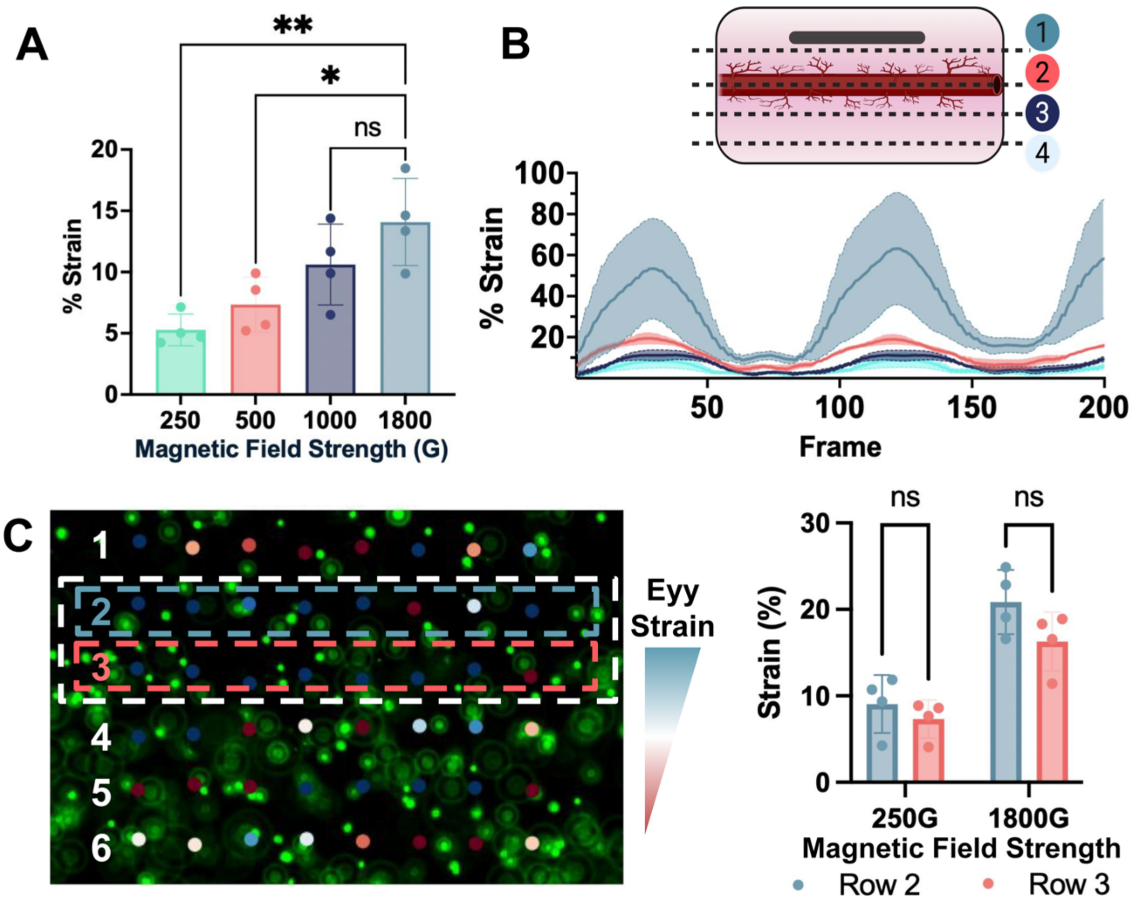


**Fig. S2.**
Spatial mapping of magnetically induced mechanical strain in the vessel-on-a-chip platform. (*A*) Magnetic field strength modulates mechanical strain at the vessel wall. Quantified strain magnitudes at 250, 500, 1,000, and 1,800 G demonstrate precise and tunable mechanical stimulation. The data are shown as mean ± SD ($^{ns}p > 0.05$, $^{*}p < 0.05$, $^{**}p < 0.01$, one-way ANOVA, n=4 independent devices). (*B*) Row-by-row spatial mapping of mechanical strain in the vessel-on-a-chip device. Top: Schematic showing measurement regions across the device, including areas proximal to the magnet (Row 1), the vessel channel (Row 2), and adjacent regions (Rows 3 and 4). Bottom: Experimental traces of X-axis strain (%) over time (frames) for each row. Shaded areas indicate the standard deviation across repeated measurements (n = 4 independent devices). (*C)* High-resolution strain mapping at the vessel boundary. Green fluorescent microspheres embedded within the collagen matrix were tracked to compute local strain (Eyy), visualized as a color-coded matrix overlaid on the construct. Left: Spatial strain map across six lateral rows. The white box outlines the vessel boundary. Row 2 (blue) corresponds to the actuator-facing side of the vessel wall, and Row 3 (red) corresponds to the opposing side. Right: Quantification of strain magnitude in Rows 2 and 3 at magnetic field strengths of 250G and 1,800G demonstrates no significant difference between actuator-facing and opposing sides. Data are shown as mean ± SD (two-way ANOVA; n = 4 independent devices; $^{ns}p > 0.05$).

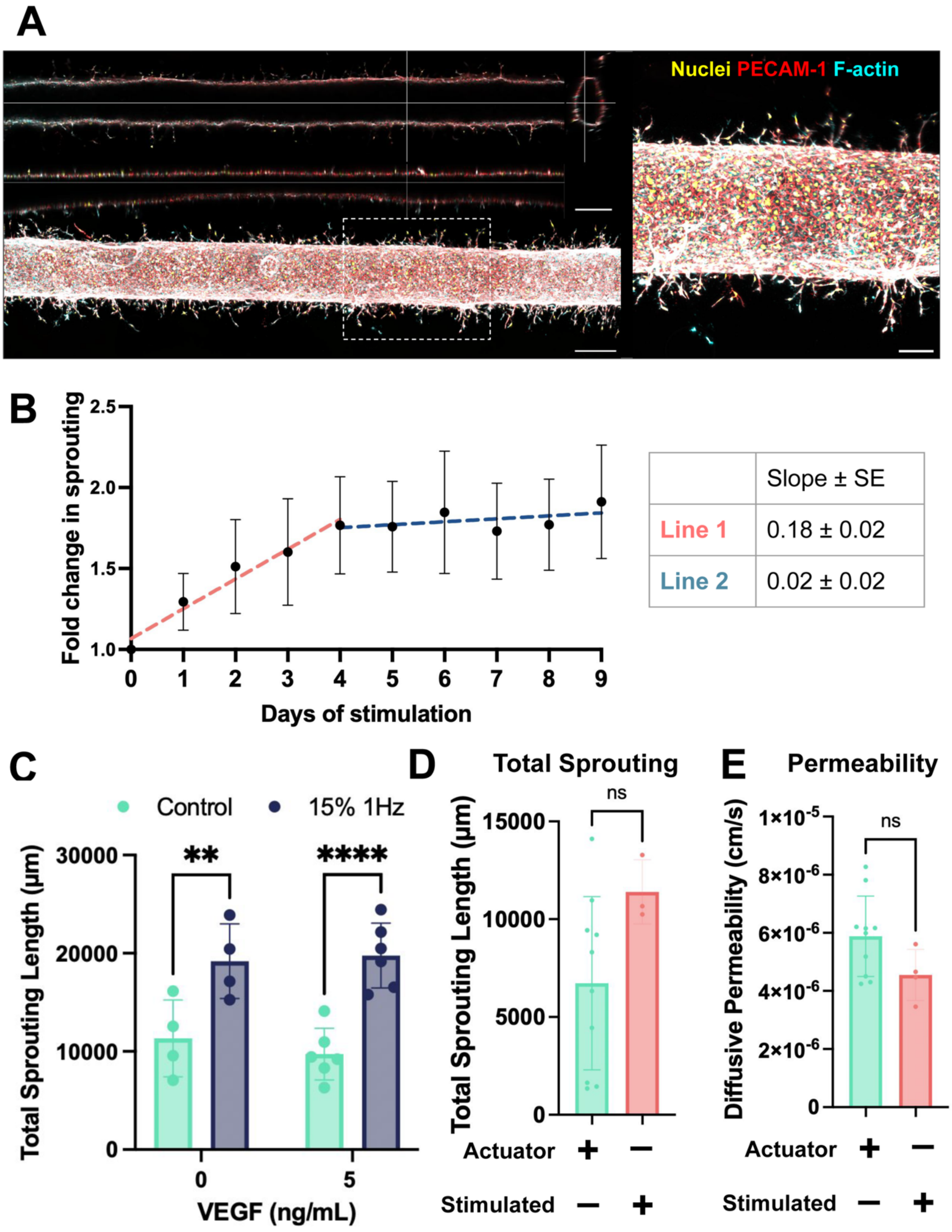


**Fig. S3.** Establishing and validating dynamic mechanical stimulation parameters for angiogenic studies. (*A*) Representative maximum intensity and projections of confocal z-stacks showing 3D sprouting from the parent vessel under 15% strain at 1Hz. Vessels stained with nuclei (yellow), PECAM-1 (red), and F-actin (cyan). Left panel shows full field view (scale bar is 500 µm); right panel shows high-magnification detail of sprouting zone with dashed box indicating region of interest (scale bar is 200 µm). (*B*) Quantification of total sprouting over 9 days of dynamic stimulation (15% at 1 Hz). Fold change in total sprouting increased during the first 4 days and plateaued thereafter, indicating saturation of the angiogenic response with prolonged stimulation. Linear regression analysis identified an early growth phase (red, slope = 0.18 ± 0.02) followed by a plateau phase (blue, slope = 0.02 ± 0.02). The data are shown as mean ± SD (n≥3 independent devices, quantified across the entire device). (*C*) Quantification of total sprouting length in vessels cultured in 0 or 5 ng/mL VEGF and subjected to 15% 1 Hz dynamic stimulation. Exogenous VEGF did not significantly affect sprouting, whereas dynamic mechanical stimulation increased angiogenesis, demonstrating a mechanically driven sprouting response. ($^{ns}p > 0.05$, *p < 0.05, **p < 0.01, ***p < 0.001, ****p < 0.0001, two-way ANOVA, n≥4 independent devices, quantified across the entire device) (*D*) Comparison of total sprouting length from parent vessel with and without the actuator magnet (Actuator), with and without permanent magnet movement (Stimulated). No significant difference in total sprouting length is observed between different control conditions. The data are shown as mean ± SD ($^{ns}p > 0.05$, Unpaired t-test, n≥3 independent devices, quantified across the entire device). (*E*) Endothelial barrier permeability assessed by diffusive permeability coefficient of 10 kDa dextran tracer. No significant difference in permeabity is observed between different control conditions. The data are shown as mean ± SD ($^{ns}p > 0.05$, Unpaired t-test, n≥4 independent devices, quantified across the entire device).

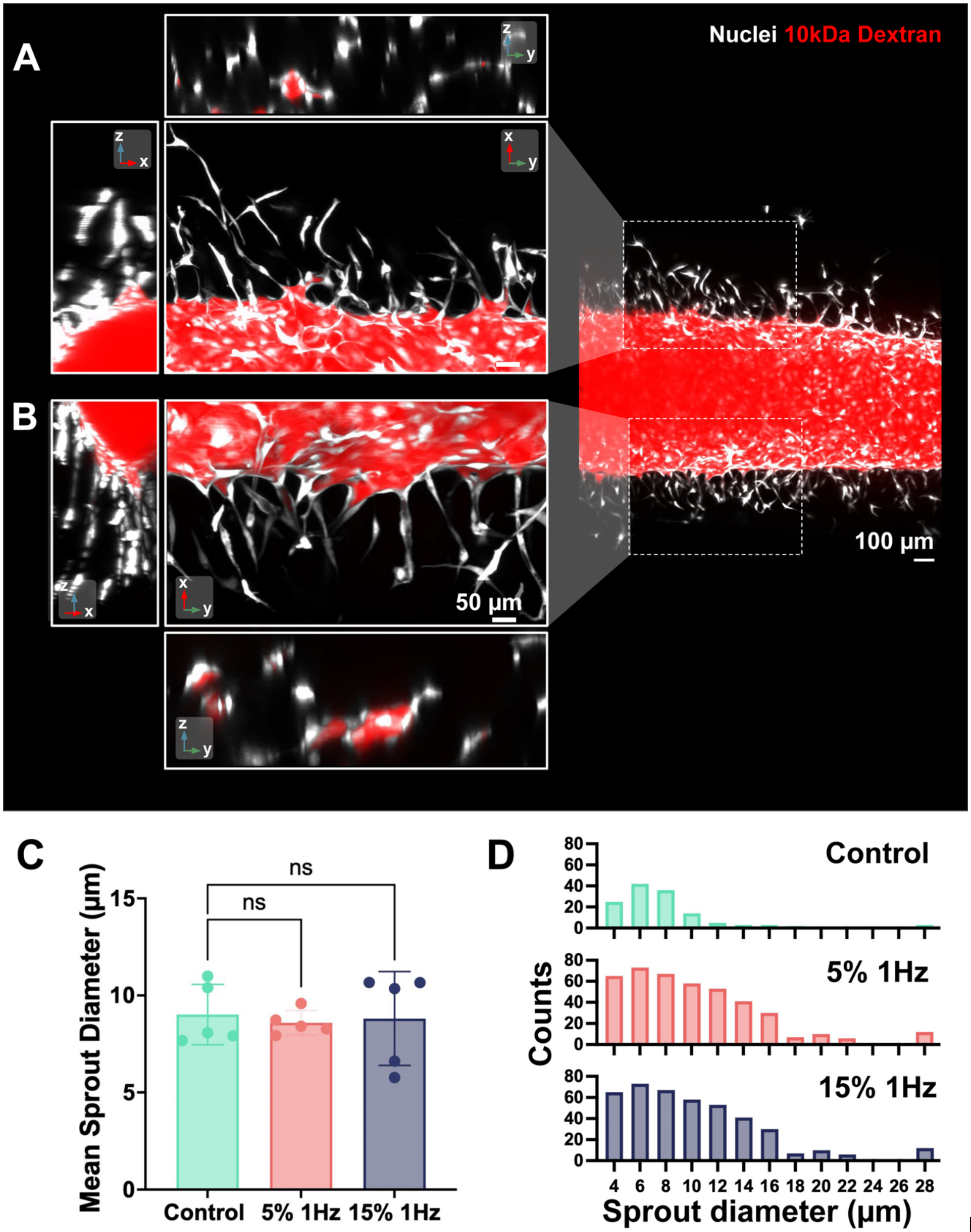


**Fig. S4.** Dynamic mechanical stimulation yields capillary-scale perfusable sprouts (A & B) Representative confocal images of multicellular sprouts perfused with 10 kDa fluorescent dextran (red). Orthogonal projections confirm luminal continuity and tracer confinement within hollow angiogenic branches. Scale bars, 100 and 50 µm.(*C*) Mean diameter of angiogenic sprouts under control, 5% 1Hz, and 15% 1Hz strain conditions. ($^{ns}p > 0.05$, one-way ANOVA, n=5 independent devices, one ROI per device) (*D*) Representative histogram distributions of sprout diameters from a single region of interest per condition (one independent device each) under control, 5% 1Hz, and 15% 1Hz strain conditions.

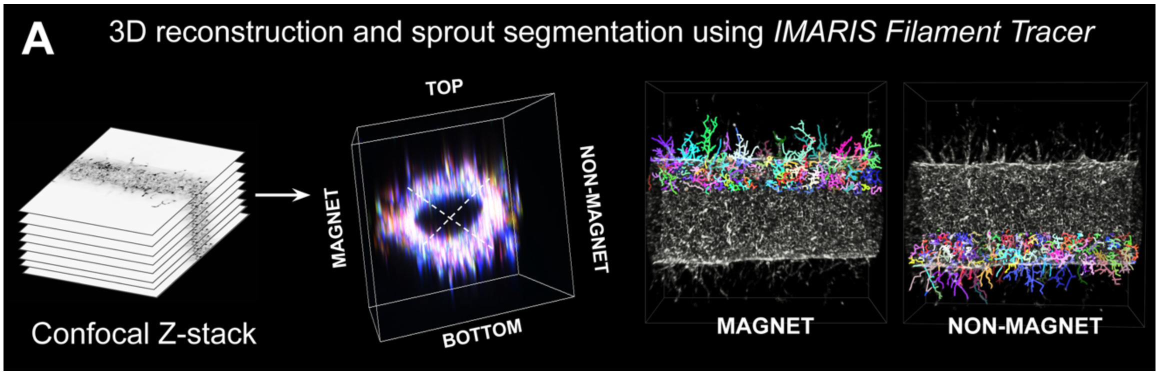


**Fig. S5.** (*A*) Schematic of the IMARIS filament tracing workflow used to reconstruct confocal z-stacks in 3D and segment angiogenic sprouts across defined vessel (top, bottom, magnet, non-magnet).

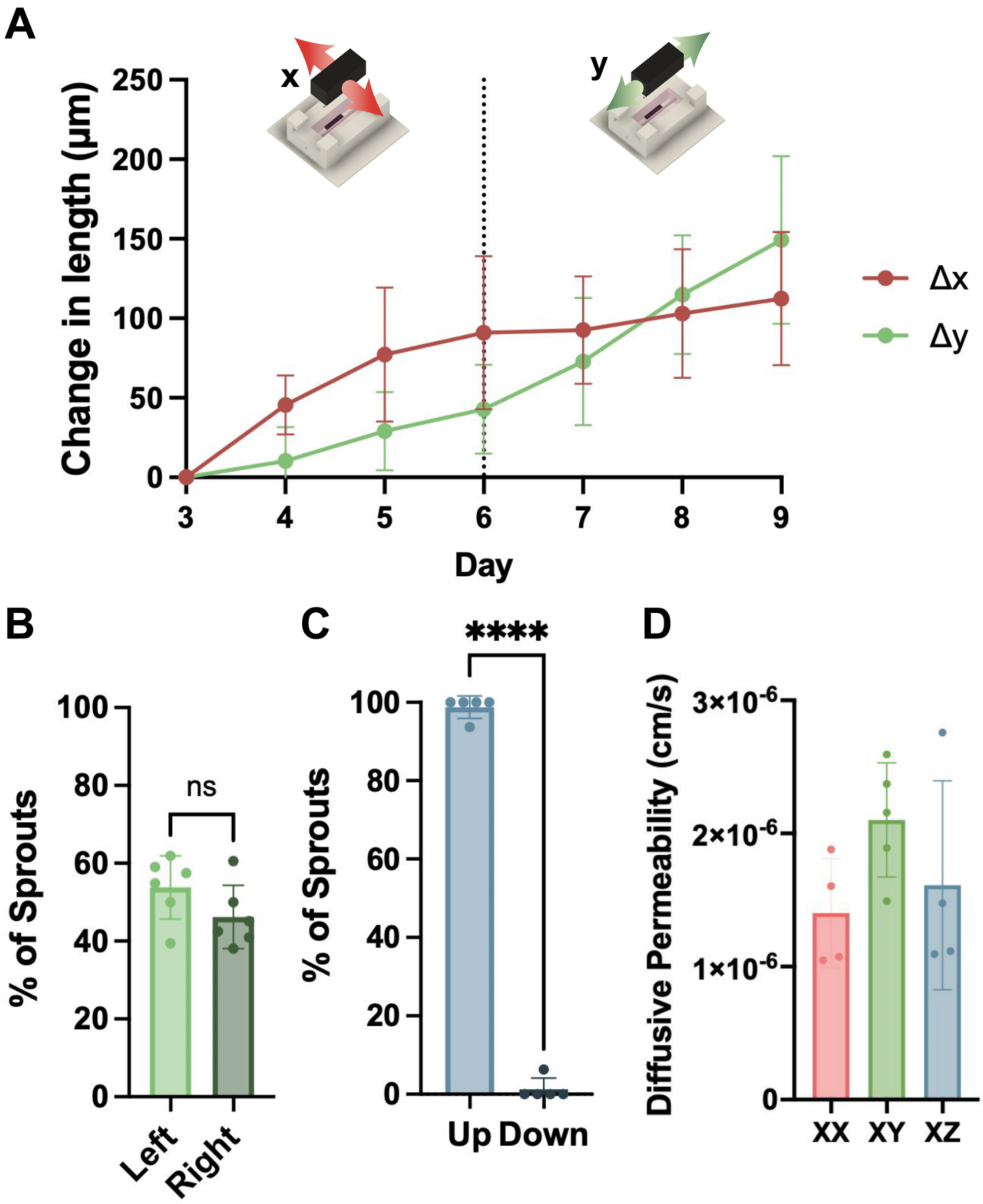


**Fig. S6.**

Quantification of sprout reorientation during sequential multi-axis stimulation. (*A*) Longitudinal changes in x- (Δx, green) and y-displacement (Δy, red) of tracked sprouts during sequential X (days 3–6) followed by Y (days 6–9) stimulation. Δx increases during X and plateaus after strain reorientation, while Δy increases following Y stimulation. (n = 5 independent sprouts, three independent devices) (B) Directionality of lateral deflection among reoriented sprouts under XY stimulation (left vs right). ($^{ns}p > 0.05$, Unpaired t-test, n≥6 independent devices, one ROI per device) (C) Directionality of vertical deflection among reoriented sprouts under XZ stimulation (up vs down) ($^{ns}p > 0.05$, $^{*}p < 0.05$, $^{**}p < 0.01$, $^{***}p < 0.001$, $^{****}p < 0.0001$, one-way

ANOVA, n≥5 independent devices, one ROI per device) (*D*) Diffusive permeability (cm/s) of vessels under XX, XY, and XZ stimulation measured using 10 kDa fluorescent dextran ($^{ns}p > 0.05$, one-way ANOVA, n≥4 independent devices, quantified across the entire device).

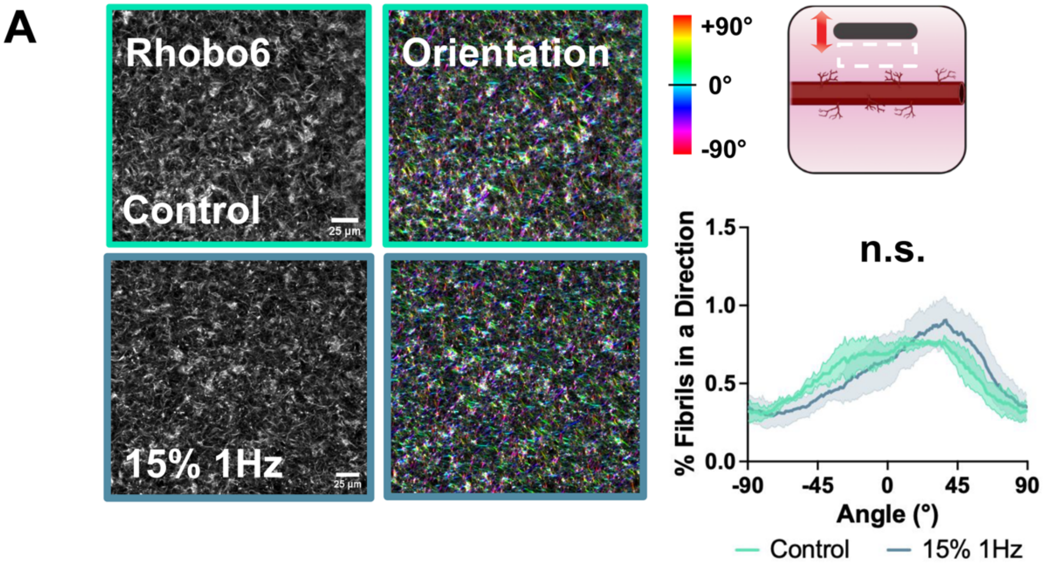


**Fig. S7.** (*A*) Collagen fibers visualized using the Rhobo6 fluorescent ECM-binding probe under unstimulated (0% strain) and dynamically actuated (15% 1Hz) conditions. Quantification of collagen fiber orientation was performed across 5 regions of interest per group ($^{ns}p > 0.05$, Unpaired t-test, n=5 ROIs).

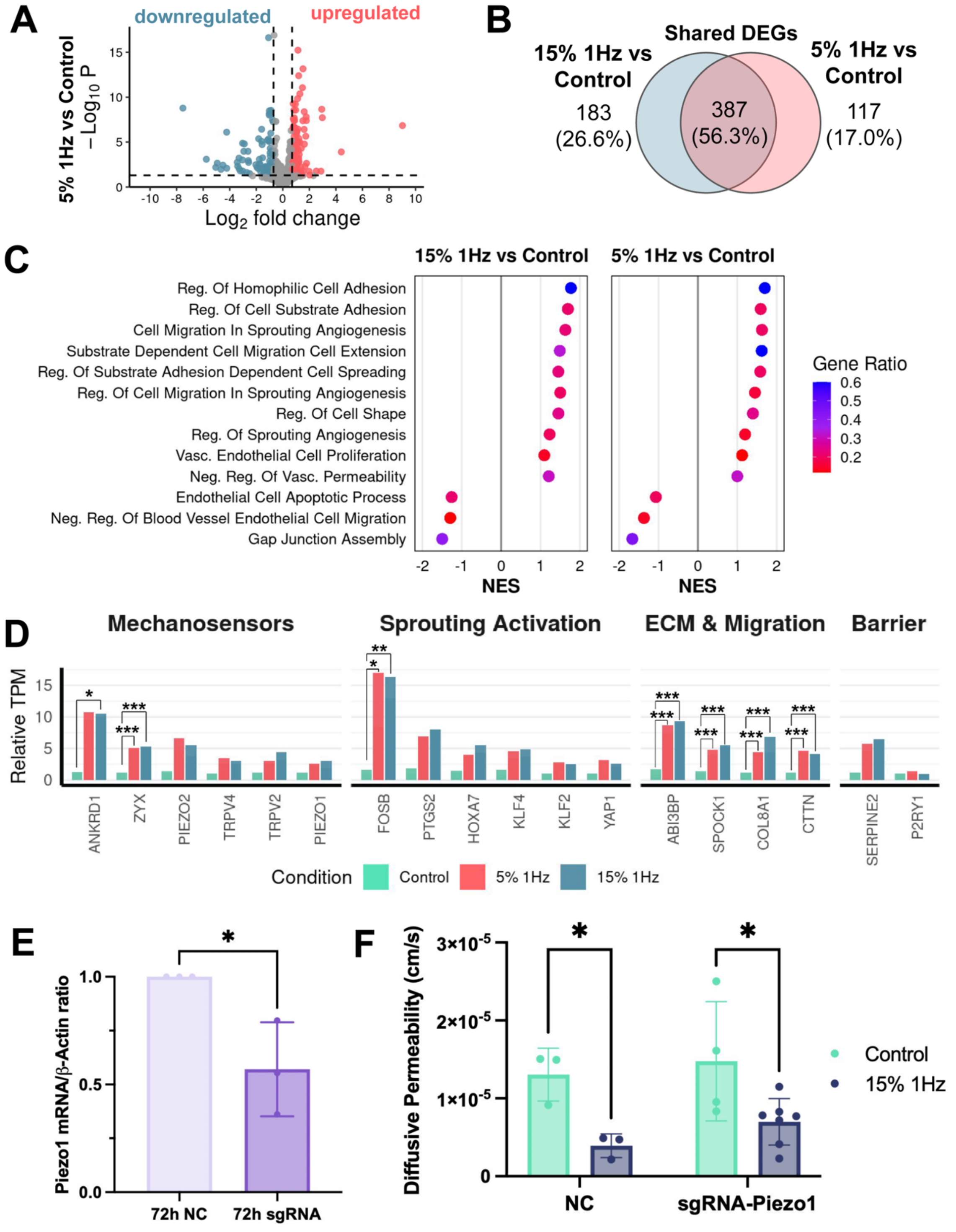


**Fig. S8.** Gene expression profiling and functional validation of mechanosensitive signaling under dynamic strain. (*A*) Volcano plots showing upregulated (red) and downregulated (blue) differentially expressed genes (adjusted $p < 0.05$, $|\log_2 \text{ fold change}| > 0.7$) for 5% 1Hz relative to control. (*B*) Venn diagrams showing the overlap of differentially expressed genes between 5% 1Hz vs control and 15% 1Hz vs control comparisons. (*C*) Gene set enrichment analysis (GSEA) of biological processes across 5% and 15% strain conditions, ranked by normalized enriched score (NES). (*D*) Relative transcripts per million (TPM) of selected genes grouped by functional category, including mechanosensors, spouting activation genes, ECM and migration-associated genes, barrier-associated genes. Significance was determined using the Wald test in DESeq2 with Benjamini–Hochberg correction for multiple hypothesis testing (FDR $\leq 0.05$). (*E*) qRT-PCR analysis of Piezo1 mRNA expression in HUVECs 72 hours post-nucleofection. Expression was normalized to a β-actin reference gene and calculated using the ΔΔCt method. ($^{ns}p > 0.05$, $^{*}p < 0.05$, two-sided Student's *t*-test, n=3 biological replicates) (*F*) Diffusive permeability (cm/s) of negative control (NC) and Piezo1 functional knockdown (sgRNA-Piezo1) vessels under control and 15% 1Hz stimulation measured using 10 kDa fluorescent dextran ($^{ns}p > 0.05$, two-way ANOVA, n≥3 independent devices, quantified across the entire device).

Movie S1 (separate file). Animated schematic of fabrication and magnetic actuation of 3D human vessel-on-a-chip platform.

Movie S2 (separate file). Fluorescent bead tracking of collagen matrix displacement under 15% 1 Hz stimulation. The relative position of the parent vessel is indicated to illustrate displacement within the surrounding matrix.

Movie S3 (separate file). Time-lapse maximum-intensity projection fluorescence sequence showing BFP-labeled endothelial nuclei imaged daily over 6 days of dynamic stimulation, corresponding to Fig. S2D.

Movie S4 (separate file). Confocal z-stack video scanning through optical sections along the z-axis of a perfused engineered vessel under 15% 1Hz dynamic strain. Red signal shows fluorescent 10 kDa dextran and white signal shows BFP-tagged nuclei. Fluorescent dextran is visible within the parent vessel lumen and extending into connected sprout lumens, demonstrating continuous, perfusable vascular structures.

Movie S5 (separate file). Three-dimensional reconstruction of a perfused engineered vessel under 15% 1 Hz dynamic strain, generated using Imaris imaging software. Red signal indicates fluorescent 10 kDa dextran and white signal indicates BFP-tagged endothelial nuclei. The reconstructed volume shows dextran confined within the parent vessel lumen and extending through the interior of 3D sprouts, demonstrating lumen continuity and perfusion under dynamic mechanical stimulation.

Movie S6 (separate file). Three-dimensional reconstruction of a single endothelial sprout extending from the parent vessel, generated using Imaris imaging software. Yellow indicates nuclei, red indicates PECAM-1, and cyan indicates F-actin. The reconstructed volume (red) reveals a hollow, continuous lumen running through the interior of the 3D sprout.

Movie S7 (separate file). Confocal maximum intensity projections of two representative sprouts imaged daily during sequential stimulation (days 3–6, x-axis strain; days 6–9, y-axis strain), corresponding to the longitudinal displacement analysis in Fig. S4A.
